# Stimulant medications affect arousal and reward, not attention

**DOI:** 10.1101/2025.05.19.654915

**Authors:** Benjamin P. Kay, Muriah D. Wheelock, Joshua S. Siegel, Ryan Raut, Roselyne J. Chauvin, Athanasia Metoki, Aishwarya Rajesh, Andrew Eck, Jim Pollaro, Anxu Wang, Vahdeta Suljic, Babatunde Adeyemo, Noah J. Baden, Kristen M. Scheidter, Julia Monk, Nadeshka Ramirez-Perez, Samuel R. Krimmel, Russel T. Shinohara, Brenden Tervo-Clemmens, Robert J. M. Hermosillo, Steven M. Nelson, Timothy J. Hendrickson, Thomas Madison, Lucille A. Moore, Óscar Miranda-Domínguez, Anita Randolph, Eric Feczko, Jarod L. Roland, Ginger E. Nichol, Timothy O. Laumann, Scott A. Marek, Evan M. Gordon, Marcus E. Raichle, Deanna M. Barch, Damien A. Fair, Nico U.F. Dosenbach

## Abstract

Prescription stimulants such as methylphenidate are being used by an increasing portion of the population, primarily children. These potent norepinephrine and dopamine reuptake inhibitors promote wakefulness, suppress appetite, enhance physical performance, and are purported to increase attentional abilities. Prior functional magnetic resonance imaging (fMRI) studies have yielded conflicting results about the effects of stimulants on the brain’s attention, action/motor, and salience regions that are difficult to reconcile with their proposed attentional effects. Here, we utilized resting-state fMRI (rs-fMRI) data from the large Adolescent Brain Cognitive Development (ABCD) Study to understand the effects of stimulants on brain functional connectivity (FC) in children (*n* = 11,875; 8-11 years old) using network level analysis (NLA). We validated these brain-wide association study (BWAS) findings in a controlled, precision imaging drug trial (PIDT) with highly-sampled (165-210 minutes) healthy adults receiving high-dose methylphenidate (Ritalin, 40 mg). In both studies, stimulants were associated with altered FC in action and motor regions, matching patterns of norepinephrine transporter expression. Connectivity was also changed in the salience (SAL) and parietal memory networks (PMN), which are important for reward-motivated learning and closely linked to dopamine, but not the brain’s attention systems (e.g. dorsal attention network, DAN). Stimulant-related differences in FC closely matched the rs-fMRI pattern of getting enough sleep, as well as EEG- and respiration-derived brain maps of arousal. Taking stimulants rescued the effects of sleep deprivation on brain connectivity and school grades. The combined noradrenergic and dopaminergic effects of stimulants may drive brain organization towards a more wakeful and rewarded configuration, explaining improved task effort and persistence without direct effects on attention networks.

## Introduction

Methylphenidate, lisdexamfetamine, and other prescription stimulants are thought to be potent, wakefulness- and attention-promoting^1^ norepinephrine and dopamine reuptake inhibitors^2,3^ used by 6.1% of Americans across all ages (and up to 24.6% of boys ages 10-19 years)^4,5^ for attention deficit hyperactivity disorder (ADHD),^6^ traumatic brain injury (TBI),^7^ narcolepsy,^8,9^ depression,^10^ as appetite suppressants,^11–14^, cognitive enhancers (nootropics),^15–17^ drugs of abuse,^18^ and to enhance athletic performance.^19^ The wakefulness-promoting properties of amphetamine were discovered in 1929, and it was later prescribed for narcolepsy and used by soldiers in World War II.^1^ Charles Bradley discovered that amphetamine seemed to treat what he termed “behavioral problem children” in 1937,^20^ although stimulants were not widely prescribed for behavior until the 1970s, and the term ADHD was not widely used until 1980.^6^ Bradley proposed that stimulants might act on attention and impulsivity by enhancing the activity of attention-promoting brain regions to increase voluntary control over action.^20^ Early research identified regions in prefrontal cortex associated with voluntary allocation of attention as being modulated by stimulants through frontostriatal circuits,^21^ while current understanding has evolved to include more diverse brain systems including sensorimotor and salience regions that serve a faciliatory role in attention.^22^ The relative effect of stimulants on these brain systems remains unclear.

The belief that stimulants act primarily on prefrontal cortex, along with evidence of beneficial effects on tasks involving attention and working memory in rodents,^23–25^ primates,^26^ and humans,^27–30^ led to the popular belief that stimulants improve attentional ability or perhaps even cognitive ability in general.^15–17^ However, closer examination of behavioral experiments shows that performance follows a U-shaped inverted curve.^25,31,32^ Lower performers improve the most with stimulants while high-performers do not improve,^23,25,26,29,30^ or even perform worse,^33^ but mistakenly perceive of their performance as improved.^34^ The most consistent behavioral effects of stimulants are improved reaction time,^24,29^ time discrimination,^29^ premature responses/impulsivity,^25,26^ effort,^33^ persistence,^27,28^ and motivation.^30,35^

Task-based fMRI studies have shown stimulant effects in prefrontal cortex as well as disparate brain regions whose functions are difficult to reconcile, e.g. insula, supplemental motor area, and thalamus.^29^ One challenge in interpreting task-fMRI results in the context of stimulants is that brain activity selectively evoked by the task contrast can be confounded by stimulant-driven differences in task performance.^36^ Resting-state fMRI (rs-fMRI) functional connectivity (FC)^37^ is not subject to performance confounds and provides a conceptual framework for synthesizing regional results into network-level hypotheses.^38–41^ While a growing number of studies have leveraged rs-fMRI to study the neural correlates of stimulants, no coherent mechanistic hypothesis of medication effects has emerged.^42^

Many prior human rs-fMRI studies,^43–48^ and some functional connectivity studies of task-fMRI data,^49^ reported significant FC changes associated with stimulants in the dorsal and ventral attention networks (DAN, VAN),^50–53^ and in cognitive control networks, such as the frontoparietal network (FPN)^54–57^ and default mode network (DMN),^58,59^ which intersect prefrontal cortex. However, these findings of stimulant-related changes in attention and control networks were not replicated in larger studies.^60,61^ Some rs-fMRI^43–45,62,63^ and positron emission tomography (PET)^63,64^ studies noted changes in primary motor regions associated with stimulants that “may be unexpected given the traditional view of ADHD as primarily involving executive control regions and networks.”^62^ Other studies^65–67^ reported stimulants affecting FC of the salience network (SAL), which is thought to govern reward- and aversion-motivated behavior.^68–70^ In several studies,^48,49,65,66,71,72^ the reported default mode or salience regions may have included portions of the parietal memory network (PMN), which is closely related to SAL^73,74^ by shared dopaminergic connections^75–77^ with the nucleus accumbens,^30,78^ and provides memory for goal-directed actions.^79–81^ Thus, stimulants may modulate cognition through multiple brain mechanisms, the relative roles of which remain incompletely understood^42^ with conflicting results from human neuroimaging studies.^43–48,60–62,64–67,71,72,82,83^

Prior human neuroimaging studies have involved sample sizes from *n* = 10^64^ to *n* = 99^60^ participants taking stimulants with relatively brief (6^46,62^ to 24 minutes^84^) fMRI acquisitions subject to reliability concerns,^85–89^ and few attempted to replicate their results in independent data or with complementary designs.^63,66^ Recent work has shown more reliable results are achieved with thousands of participants for brain-wide association studies (BWAS),^90^ extended duration, repeated fMRI scans for precision functional mapping (PFM) studies,^86,88,91–94^ and the use of discovery and replication sets for validation.^91,92,95^ Many prior analyses used a region of interest (ROI) approach^43–48,60,64–67^, whereas advances in computational methods have now enabled data-driven approaches with increased statistical power.^96–98^ Prior imaging studies did not control for the effects of sleep, even though inadequate sleep (less than 9 hours of sleep per night in children)^99^ is common^100,101^ and associated with cognitive decrements.^102^

The unexpected^62^ relationship between stimulants and primary motor cortex has been interpreted as relating to inhibition of motoric output in hyperactive individuals^21,82^ based on the observation that ADHD is associated with decreased primary motor cortex short interval cortical inhibition in behavioral^103^, fMRI^104^, and transcranial magnetic stimulation (TMS) studies.^82,105–108^ However, recent findings in functional neuroanatomy and connectomics provide an alternative context in which to interpret stimulant-related differences in motor cortex. Multi-modal precision imaging research has shown that primary motor cortex is not a simple homunculus, but is interleaved with the somato-cognitive action network (SCAN)^109^ with diverse functions including regulation of sympathetic outflow.^110^ Action^111^ and motor regions reflect arousal state^112–114^ such that FC within motor (SM), auditory (AUD), and visual (VIS) networks is increased during sleep and decreased during wakefulness.^86,115–118^

In this study we used rs-fMRI data from the Adolescent Brain Cognitive Development (ABCD) Study (*n* = 11,875).^119,120^ We employed a data-driven whole-connectome strategy to model differences in FC in attention, arousal, and salience/memory networks related to prescription stimulants without a priori exclusion of other networks. Network level analysis (NLA)^121,122^ was used to account for multiple comparisons. The findings were validated^91,92^ with a precision imaging drug trial (PIDT) of methylphenidate 40 mg in healthy adults without ADHD (165-210, mean 186 minutes of rs-fMRI data per participant).^123^

## Results

### Stimulant use is prevalent in children

In the ABCD Study (*n* = 11,875, 8-11 years old, data collected 2016-2019), 7.8% of children (74.6% boys) were prescribed a stimulant and 6.2% (74.0% boys) took the stimulant on the morning of their MRI scans. See Supplemental Figure 6 for a breakdown of stimulants by active ingredient. Data on specific dose and formulation were not available in the ABCD Study. Using stringent criteria,^124^ 3.7% of children (69.4% boys) had ADHD of whom 42.7% were prescribed a stimulant and 34.9% took the stimulant on the morning of scanning. Only 20.7% of children who took a stimulant on the morning of scanning met criteria for ADHD. Using less stringent criteria for identifying ADHD (see Methods) 76.2% of children taking a stimulant had ADHD. A sample of *n* = 5,795 children with complete data, including sufficient low-motion fMRI, included *n* = 337 (73.0% boys) children taking a stimulant on the morning of their scans.

### Stimulants change children’s action, motor and salience connectivity

To visualize the relative FC differences (*t* -values, see Methods for covariates) associated with pre-scan stimulant use in each brain region, we computed the magnitude of FC differences over the edges connected to each region. The largest stimulant-related FC differences were in somato-cognitive action, primary motor, auditory, salience, and parietal memory regions, see Figure 1(a). An exemplar parcel-wise seed map using a motor-hand parcel with the greatest FC difference is shown in Figure 1(b). For the full FC matrix see Supplemental Figure 2. The nucleus accumbens is thought to be central to dopamine-mediated processing of reward, salience, and effort.^125,126^ An additional nucleus accumbens seed map showed high FC with canonical salience regions in cortex (e.g. anterior inferior right insula)^68,70^ but no significant difference related to stimulants, see Supplemental Figure 3.

**Figure 1:**
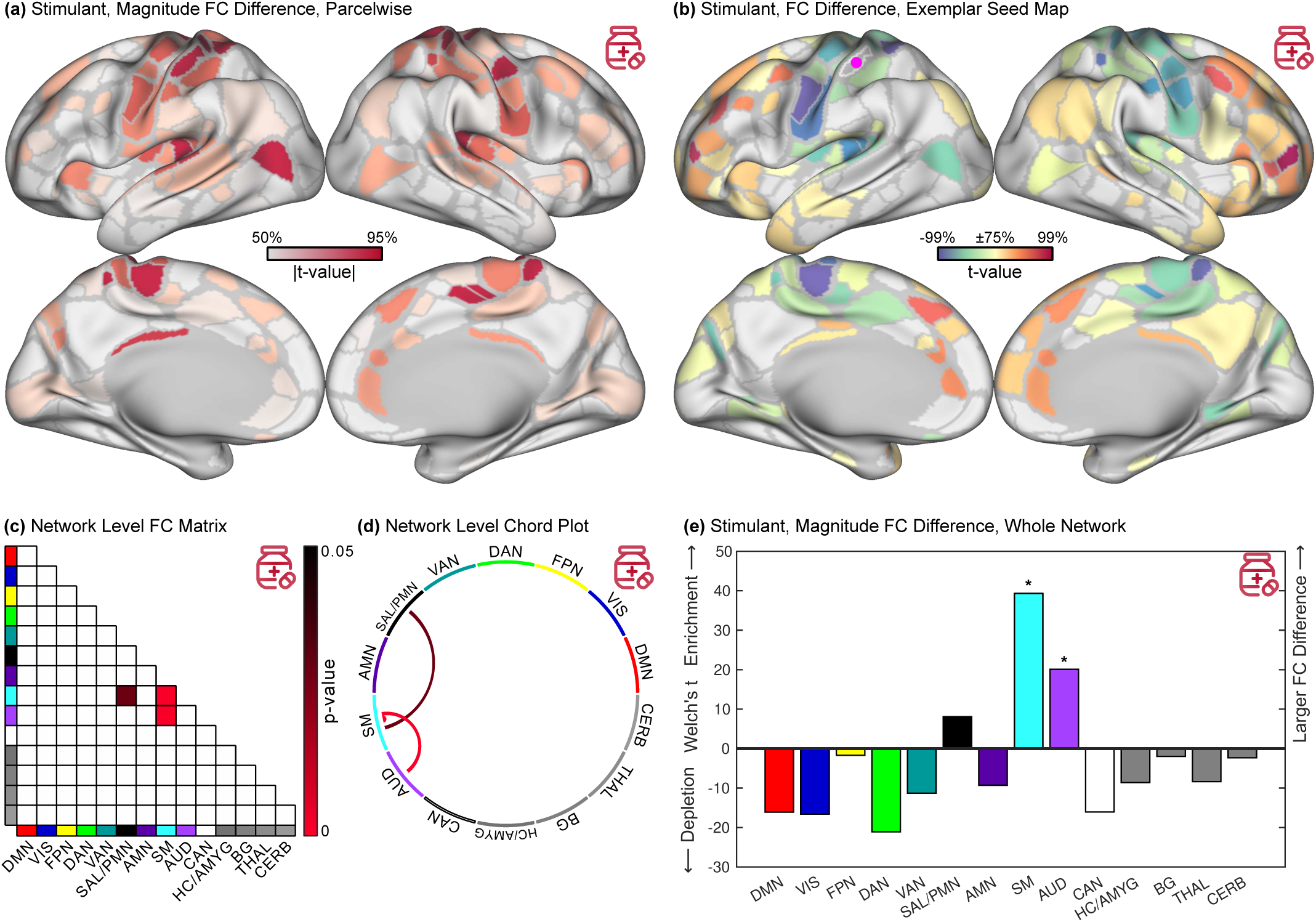
Stimulant related functional connectivity differences. ABCD Study data 5,795 children, 337 taking a stimulant. Stimulant-related findings are color coded red. **(a)** Magnitude (root mean square) of functional connectivity (FC) difference shown on the Gordon-Laumann cortical parcellation.^133^ **(b)** Differences in FC with an exemplar (most affected by stimulants) seed parcel in the motor-hand region (purple dot). **(c,d)** Significant (FWER p < 0.05) differences in FC between network pairs. **(e)** Magnitude (Welch’s *t* -statistic) of FC differences in whole networks relative to the whole connectome. Significant (FWER p < 0.05) differences are indicated by a **\***. DMN: default mode, VIS: visual, FPN: fronto-parietal, DAN: dorsal attention, VAN: ventral attention, SAL: salience, PMN: parietal memory, AMN: action-mode, SM: somato-cognitive action/motor, AUD: auditory, CAN: context association, HC: hippocampus, AMYG: amygdala, BG: basal ganglia, THAL: thalamus, CERB: cerebellum.

The somato-cognitive action (SCAN) and motor networks, which are interleaved along the central sulcus, were treated as one somato-motor (SM) network for the purpose of statistical comparison. Among pairs of canonical networks, stimulants were associated with significantly decreased FC within and between SM and auditory (AUD) networks (NLA, Westfall-Young step-down FWER-corrected^127^ *P* < 0.05). Stimulants were associated with significantly increased FC between SM and salience/parietal memory networks (SAL/PMN). See Figure 1(c,d). Among all edges within and between each network, stimulants were associated with the largest differences in FC in SM and AUD (FWER *P* < 0.05) and a trend toward relatively larger FC differences in SAL/PMN, see Figure 1(e). There were no significant FC differences in attention (DAN, VAN) or control (FPN) networks, despite 95% power to detect stimulant-related differences in attention networks, see Supplemental Table 2.

It has been hypothesized that children with ADHD may show different changes in FC in response to stimulant intervention during an attention-demanding task compared to rest.^49^ The n-back task was used in the ABCD Study to engage working memory and cognitive control in adolescents.^128^ Functional MRI data from the n-back task was treated as rest and analyzed without regressing out the task paradigm. Stimulant related differences in n-back FC were parcel-wise highly correlated with those of resting FC (*r* = 0.45, spin test^129,130^ *P* = 0.0015), see Supplemental Figure 4. Stimulants were not associated with significant differences in task-evoked fMRI activation for 0-back vs fixation, see Supplemental Figure 5, although power may have been limited by fewer children with high-quality n-back data (*n* = 109 taking stimulants) and technical issues specific to task design in the ABCD Study.^131^

Differences in the precise molecular action of different stimulant drugs have been reported.^3^ An analysis of the ABCD data separating stimulants into specific drugs (methylphenidate, lisdexamfetamine, etc.) showed the same pattern of FC differences for each drug, see Supplemental Figure 6. The stimulant-related patten of FC differences was not observed for cetirizine, a common allergy medication taken by *n* = 291 children on the day of scanning that is not psychoactive and was therefore chosen as a negative control.^132^ The cetirizine related differences in FC, which were below the threshold for significance, were parcel-wise not correlated with those of stimulant on the cortex (*r* = 0.059, spin-test *P* = 0.39), see Supplemental Figure 7.

Stimulant-related FC differences were specifically associated with taking the stimulant drug on the morning of scanning. The subset of children (*n* = 76) who were prescribed stimulants but did not take them on the morning of scanning showed no significant FC differences compared to *n* = 5,382 children not prescribed or taking a stimulant, and the 337 children taking a stimulant on the day of scanning showed the same pattern of connectivity when compared to the 76 stimulant-users who did not take their stimulant on the day of scanning as they did when compared to children who were not prescribed a stimulant, see Supplemental Figure 8. Results were not due to differences in ADHD diagnosis or head motion, see Supplemental Figures 9, 10, and Supplemental Table 3.

### Stimulant-driven connectivity changes validated in an adult trial

The ABCD Study does not experimentally control for why children take stimulants. Therefore, differences in FC associated with stimulants were validated^91,92^ in a trial with 5 healthy adult participants (165-210 minutes of rs-fMRI data each). Each participant had 120-180 minutes of rs-fMRI data off stimulants and 15-60 minutes of rs-fMRI data on methylphenidate (Ritalin) 40 mg.^123^ The study design controlled for factors correlated with stimulant use (e.g. ADHD diagnosis) by recruiting participants who were not prescribed a stimulant and comparing FC within the same individuals on- and off-stimulant. The largest stimulant-related changes in FC in these controlled data were the same as in ABCD: decreased within-network FC in SM (mixed effects *P* -value 0.008) and increased cross-network FC between SM and SAL/PMN (*P* = 0.013). Parcel-wise correlation between the two studies’ magnitude FC difference maps was *r* = 0.32 (cortex-only *r* = 0.36, spin test *P* < 0.0001), see Figure 2. For edge-wise correlation between the two studies see Supplemental Figure 11.

**Figure 2:**
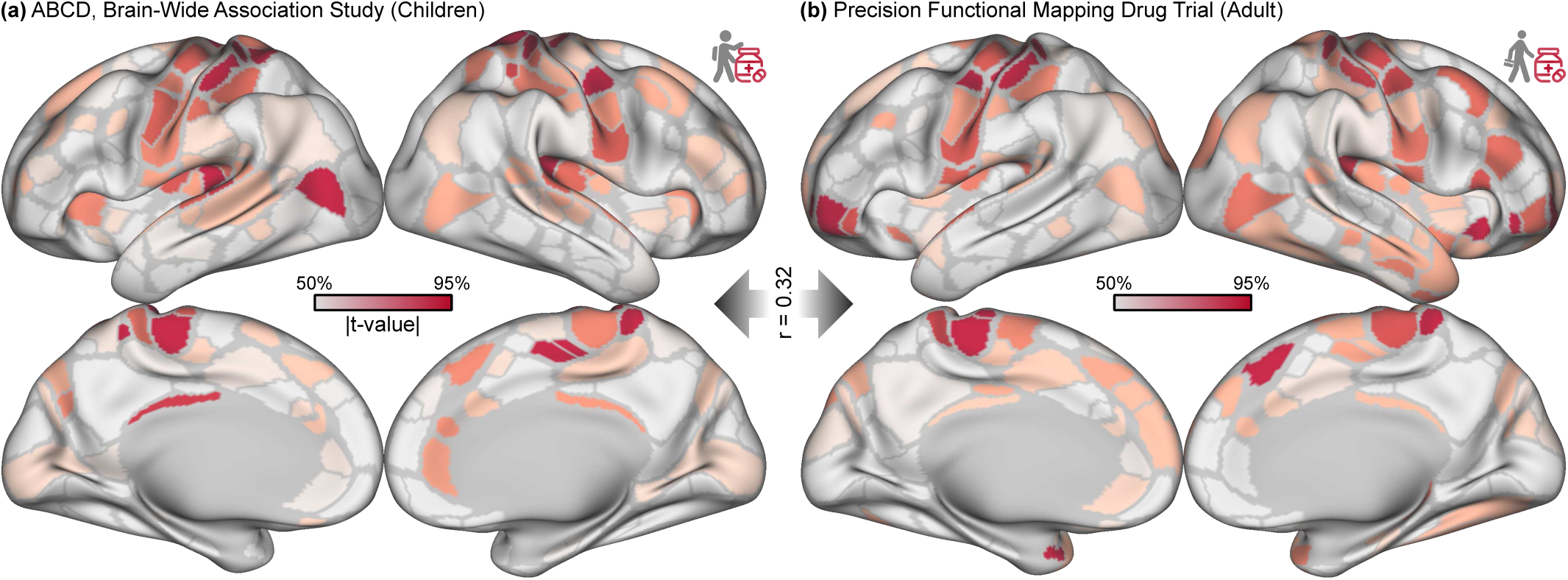
Stimulant effects validated in precision imaging drug trial. **(a)** Magnitude (root mean square) of functional connectivity (FC) differences shown on the Gordon-Laumann cortical parcellation^133^ for 337 children taking stimulants in the ABCD Study (total *n* = 5,795). **(b)** Magnitude of acute FC differences in adult participants (*n* = 5) given methylphenidate 40 mg in a controlled study. The cortical maps are correlated at *r* = 0.36 (spin-test *P* < 0.001).

### Stimulants mimicked the effects of getting more sleep

The greatest stimulant-related differences in FC were in somato-cognitive action and motor networks (SM) associated with arousal/wakefulness,^112–118^ therefore we characterized the FC pattern associated with getting more sleep and compared it to the FC pattern associated with taking stimulants. Parents of children in the ABCD Study were asked, “How many hours of sleep does your child get on most nights?” ^134^ Parent-reported sleep duration served as a surrogate measure of being better rested, or arousal/wakefulness, at the time of scanning.

Longer sleep duration was associated with FC differences in motor, auditory, and visual regions in a pattern similar (cortex+subcortex *r* = 0.58, cortex-only *r* = 0.58, spin-test *P* < 0.0001) to that of taking a stimulant, see Figure 3(a). Sleep duration was also extremely similar to stimulants in the exemplar parcel-wise seed map (cortex+subcortex *r* = 0.87, cortex-only *r* = 0.86, spin-test *P* < 0.0001), see Figure 3(b). See Supplemental Figure 2 for the full FC matrix. At the level of network pairs, sleep duration was associated with significantly (FWER *P* < 0.05) decreased FC within SM and decreased FC between SM and primary sensory networks (auditory AUD and visual VIS), see Figure 3(c,d). At the level of whole networks, sleep duration was associated with significant (FWER *P* < 0.05) changes in SM, AUD, and VIS. Thus, while stimulant- and sleep-related patterns of FC were similar, stimulants were associated with greater relative differences involving SAL/PMN than sleep duration.

**Figure 3:**
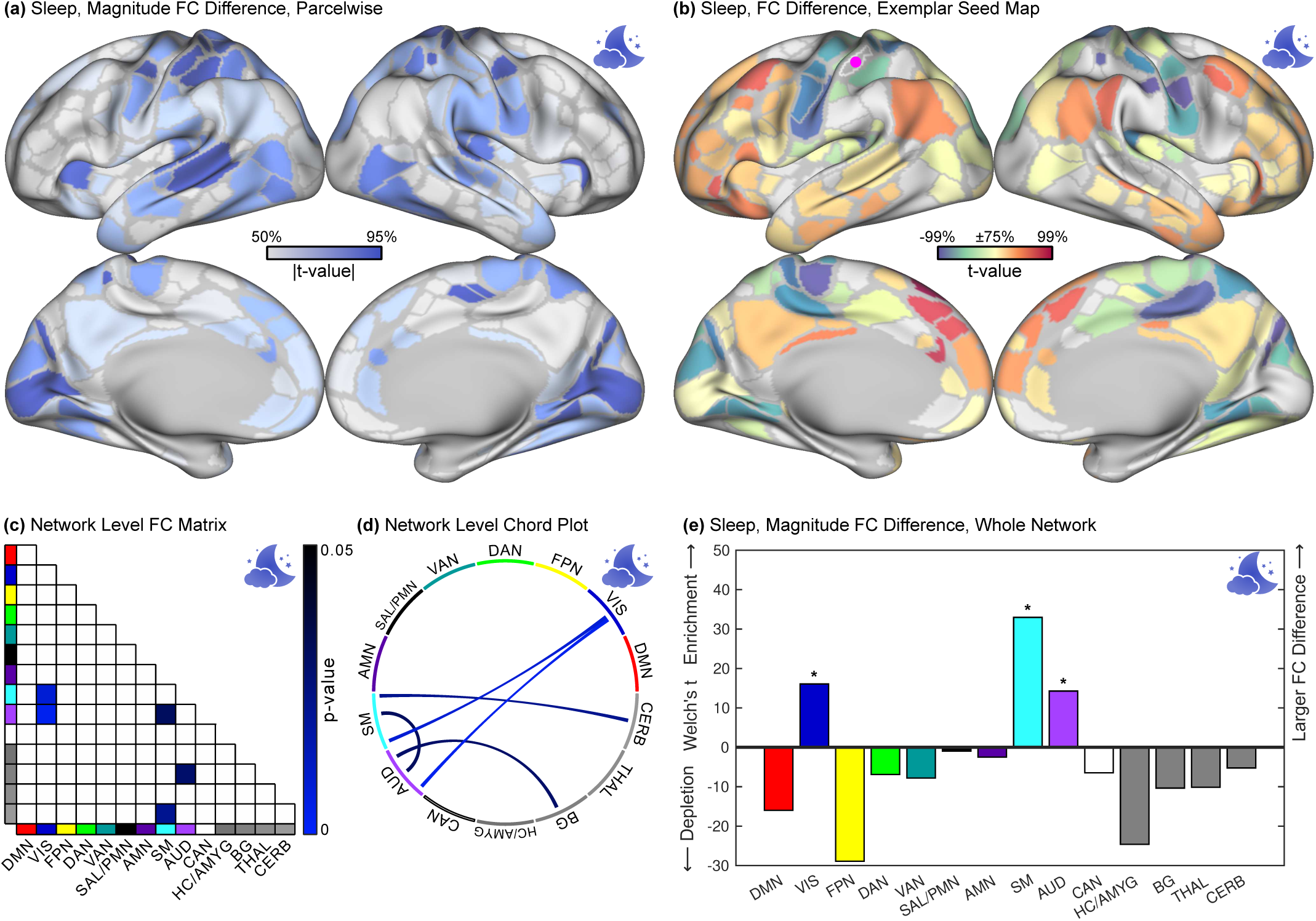
Sleep duration related functional connectivity differences. ABCD Study data 5,795 children. Sleep-related findings are color coded blue. **(a)** Magnitude (root mean square) of functional connectivity (FC) differences shown on the Gordon-Laumann cortical parcellation.^133^ **(b)** Differences in FC with an exemplar seed parcel in the somatomotor hand region (purple dot). **(c,d)** Significant (FWER *P* < 0.05) differences in FC between network pairs. **(e)** Magnitude (Welch’s *t* -statistic) of FC difference in whole networks relative to the whole connectome. Significant (FWER *P* < 0.05) changes are indicated by a **\***. DMN: default mode, VIS: visual, FPN: fronto-parietal, DAN: dorsal attention, VAN: ventral attention, SAL: salience, PMN: parietal memory, AMN: action-mode, SM: somato-cognitive action/motor, AUD: auditory, CAN: context association, HC: hippocampus, AMYG: amygdala, BG: basal ganglia, THAL: thalamus, CERB: cerebellum.

### Arousal regions showed the strongest sleep-related connectivity differences

To further validate our finding of decreased arousal-related FC within SM, AUD, and VIS, we used data from three independent studies. Sleep was correlated with an arousal template derived from correlation of EEG alpha slow wave index (alpha/delta power ratio) with fMRI signal (*n* = 10)^117,118^ at *r* = 0.49 (spin test *P* < 0.0001), see Figure 4(b) and Supplemental Table 4. Sleep was correlated with a second arousal map derived from coherence of respiratory variation with fMRI signal^113^ from *n* = 190 participants with real-time respiratory data in the Human Connectome Project^135^ at *r* = 0.51 (spin test *P* = 0.0015), see Figure 4(c).

**Figure 4:**
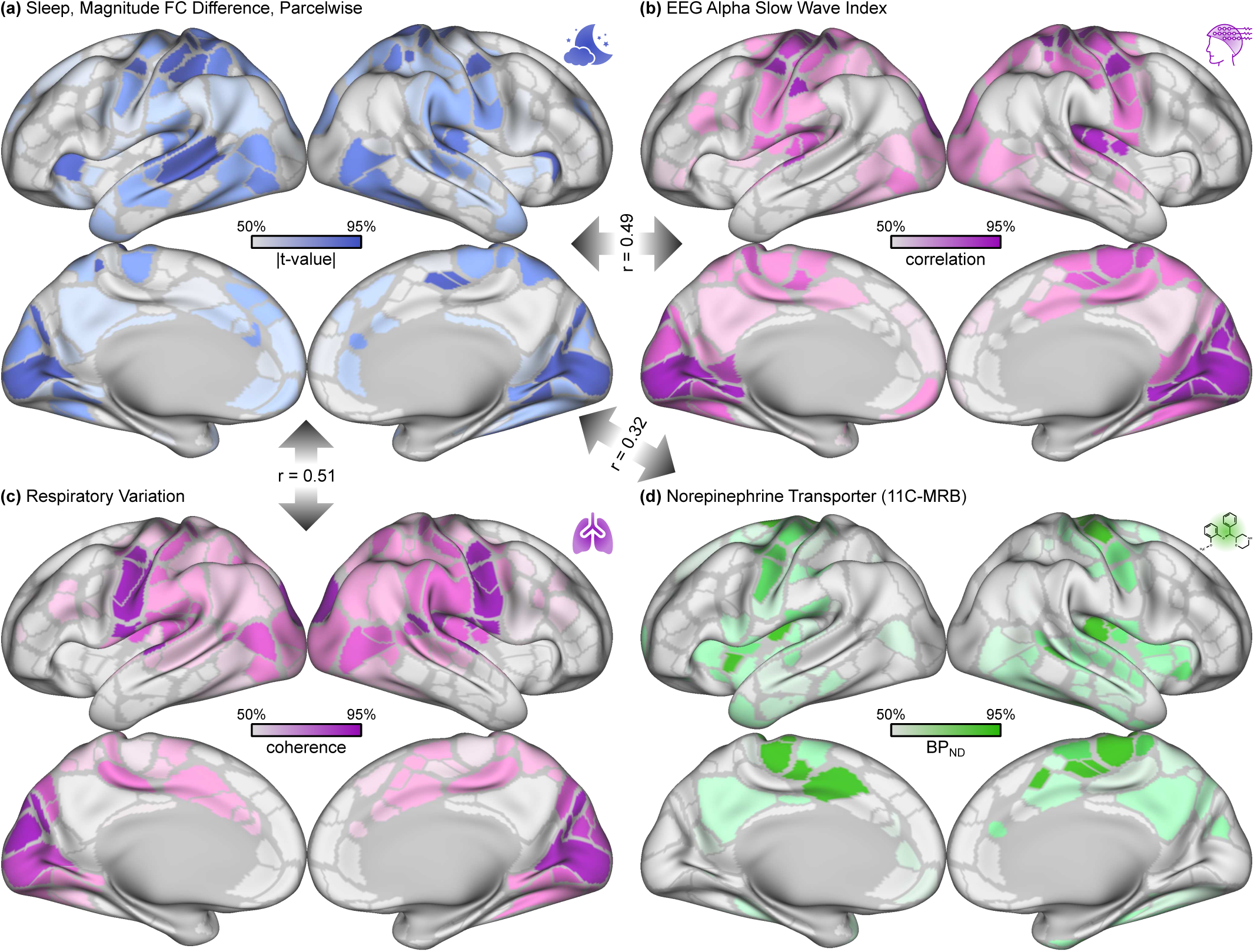
Sleep duration effects validated against independent brain maps of arousal. **(a)** Magnitude (root mean square) of functional connectivity (FC) differences related to sleep duration shown on the Gordon-Laumann cortical parcellation^133^ (ABCD Study, *n* = 5,795). **(b)** Arousal template obtained by correlating EEG alpha slow wave index (alpha/delta power ratio) with fMRI signal intensity (*n* = 10).^117,118^ **(c)** Arousal map obtained from coherence between respiratory variation and fMRI signal intensity based on (Human Connectome Project, *n* = 190).^113^ **(d)** Non-displaceable binding potential for 11C-MRB (methylreboxetine) in a positron emission tomography (PET) study (*n* = 20).^137,138^ Correlations between cortical maps are shown in gray arrows and summarized in Table 4. The correlation between the EEG- and respiration-derived arousal maps was *r* = 0.60 (spin test *P* < 0.0001).

Stimulants increase synaptic levels of norepinephrine,^2,3^ a neurotransmitter strongly associated with arousal,^136^ therefore we compared FC differences related to sleep duration with a PET map of norepinephrine transporter (NET) density (*n* = 20).^137,138^ Sleep and NET density were significantly correlated at *r* = 0.32 (spin test *P* = 0.005) in cortical parcels, see Figure 4(d) and Supplemental Table 4. Receptor density maps for dopamine, which is modulated by stimulants but less strongly associated with arousal,^136^ are shown in Supplemental Figure 13.

Stimulant-related FC differences were also significantly (spin-test *P* < 0.05) correlated with maps of arousal and norepinephrine receptor density, see Supplemental Table 4.

### Stimulants and sleep had similarly beneficial effects on performance

Stimulants^27,28^ and getting sufficient sleep^139,140^ are both thought to have beneficial effects on attention and working memory. The ABCD Study collected data on parent-reported school letter grade, out-of-scanner performance on the NIH Toolbox,^141^ and in-scanner performance on the n-back task. These cognitive measures were modeled against stimulants taken on the day of scanning and sleep duration with age, sex, and socioeconomic covariates, see Methods. ADHD was associated with significantly worse school grades, NIH Toolbox performance, and rate of correct responses on the n-back, while getting more sleep was associated with significant improvement in all of these measures, see Table 1. Children with ADHD who took a stimulant had improved cognitive performance on all measures compared to those who did not take a stimulant (significant ADHD ×stimulant interaction), and children with less sleep had better school grades if they took a stimulant (significant, negative stimulant ×sleep interaction). Children getting adequate sleep who did not have ADHD did not have better school grades, NIH Toolbox scores, or rate of correct responses on the n-back compared to those who did not take a stimulant. Taking a stimulant did significantly improve reaction time on the n-back by about 100 ms independent of other factors. Thus, overall, stimulants improved cognitive performance only for participants with ADHD or insufficient sleep (see *P* -values in Table 1).

**Table 1:**
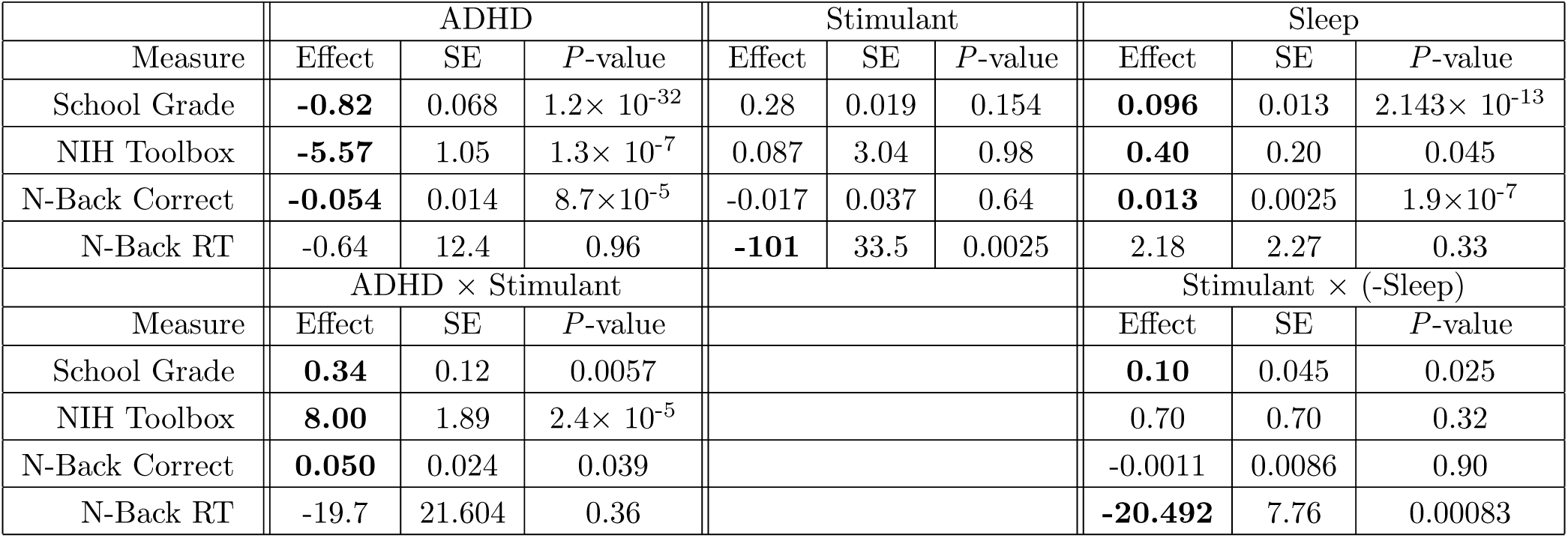
Differences in cognitive performance related to ADHD, stimulants, and sleep. A linear regression model was used to predict school letter grade (1 = F, 5 = A), NIH Toolbox score (mean 50, SD 10),^141^ n-back correct response rate (1 = 100% correct), and n-back reaction time (RT, in milliseconds) from ADHD diagnosis and sleep duration (hours) with sex, age, and socioeconomic factors as covariates in *n* = 5,795 children, 337 taking stimulants. ADHD and sleep were each associated with significant improvements on cognitive performance, while stimulants were observed to most improve performance for children with ADHD (ADHD ×stimulant interaction) or sleep deprivation (stimulant ×-sleep interaction).

### Stimulants rescued sleep-deficit induced changes

Only 48% of children in the ABCD Study were reported by their parents as getting the recommended^99^ 9 or more hours of sleep per night. Taking stimulants and longer average sleep duration (being better rested) had similar effects on brain connectivity. Therefore, we performed subanalyses of the relations of sleep to behavior and FC in subsets of children taking and not taking stimulants. There was no significant association between taking a stimulant and sleep duration, after accounting for ADHD diagnosis (*P* < 0.0001), see Supplemental Figure 12. Behaviorally, children who slept longer (per parent report) had significantly better school grades, NIH Toolbox scores, and rate of correct responses on the n-back, see Table 1. Conversely, children with less sleep had significant decrements in their cognitive performance. However, the deleterious association of sleep deprivation with cognitive performance was not significant in the subset of children taking stimulants (*n* = 337). Children getting less sleep but taking a stimulant (stimulant × -sleep interaction term) received grades that were significantly better than those of children getting less sleep not taking a stimulant, and equal to the grades of well-rested children not taking a stimulant (Table 1).

Longer sleep duration was associated with decreased within-network connectivity in somato-cognitive action/motor, auditory, and visual regions in children not taking stimulants, see Figure 5(a). Conversely, children with shorter sleep duration who were relatively sleep-deprived had increased within-network connectivity in SM, AUD, and VIS. These sleep-related differences in FC closely mirrored those in the whole cohort, see Figure 3(a). Remarkably, the relationship between sleep and FC vanished in the subset of children taking stimulants, see Figure 5(b) and Supplemental Figure 14 for the full FC matrix.

**Figure 5:**
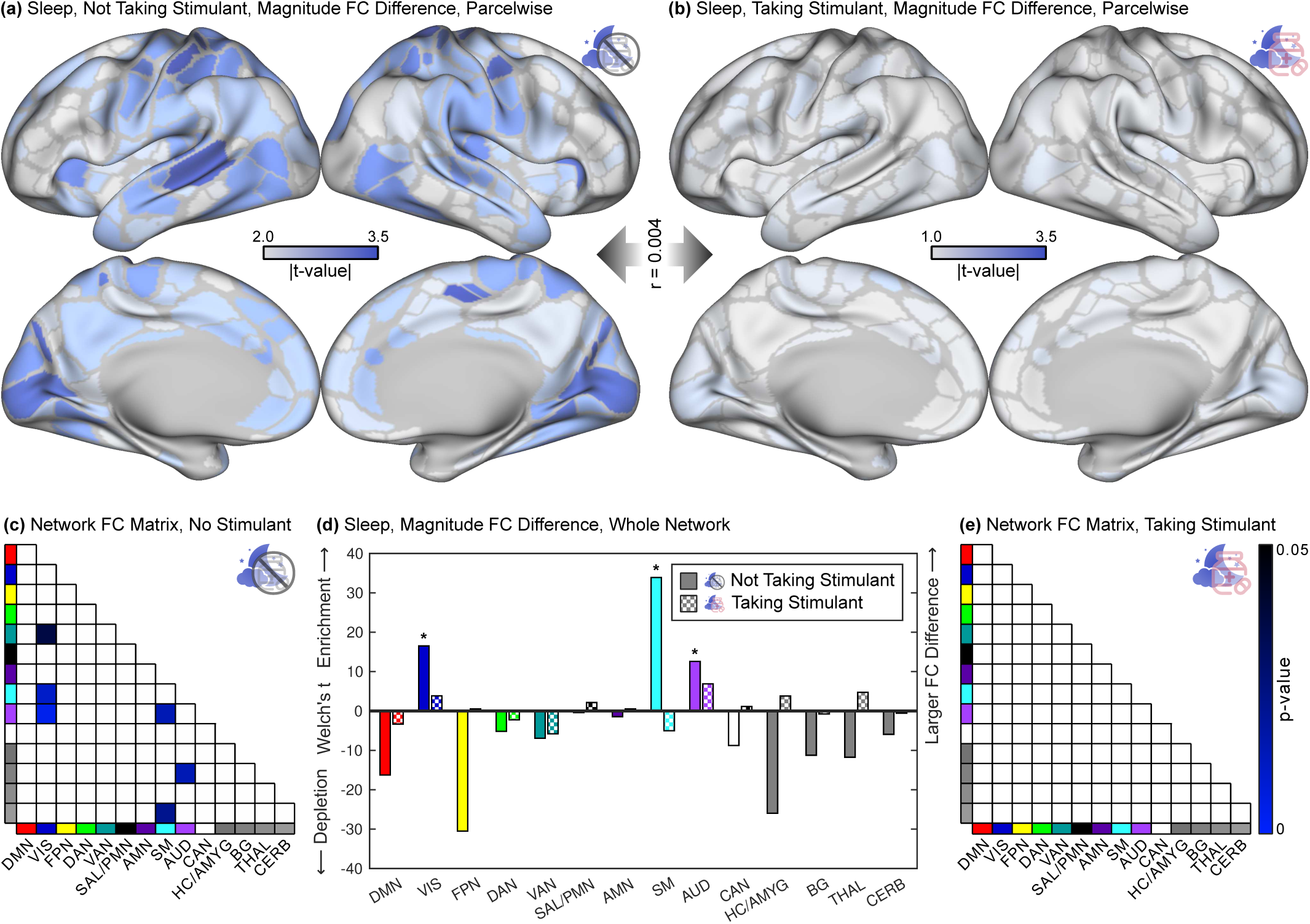
Sleep duration and stimulant use’s interacting brain effects. ABCD Study data 5,795 children, 337 taking a stimulant. **(a)** Functional connectivity (FC) difference magnitude (root mean square) for sleep shown on the Gordon-Laumann cortical parcellation^133^ in children not taking stimulants (*n* = 5,458) and **(b)** taking stimulants (*n* = 337). A more liberal *t* -value threshold was used in (b) to show detail. **(c)** Significant (FWER *P* < 0.05) differences in FC between network pairs in children not taking stimulants. **(d)** Magnitude (Welch’s *t* -statistic) of FC differences in whole networks, relative to the whole connectome, for sleep in children not taking stimulants and taking stimulants. Significant (FWER *P* < 0.05) changes are indicated by a **\***. **(e)** Significant (FWER *P* < 0.05) differences in FC between network pairs in children taking stimulants. DMN: default mode, VIS: visual, FPN: fronto-parietal, DAN: dorsal attention, VAN: ventral attention, SAL: salience, PMN: parietal memory, AMN: action-mode, SM: somato-cognitive action/motor, AUD: auditory, CAN: context association, HC: hippocampus, AMYG: amygdala, BG: basal ganglia, THAL: thalamus, CERB: cerebellum.

The pattern of sleep related FC differences were (parcelwise) very different in children taking a stimulant compared to children not taking a stimulant, *r* = -0.026 (cortex-only *r* = 0.0004, spin-test *P* = 0.997). Sleep was associated with significant (FWER *P* < 0.05) differences in SM, AUD, and VIS in children not taking stimulants, see Figure 5(c,d). There were no significant differences in FC between canonical network pairs or whole networks in the subset of children taking stimulants, see Figure 5(d,e). The edgewise sleep × stimulant interaction and the difference in sleep-related FC between children taking and not taking stimulants (Wald test)^142^ are shown in Supplemental Figure 15. The difference persisted after matching for sample size (subsampling to *n* = 337 children), see Supplemental Figure 16. The pattern of stimulant-related FC differences was more similar to the pattern of sleep-related FC differences in stimulant-takers with less sleep, see Supplemental Figures 17 and 18.

## Discussion

### Stimulants modulate arousal and salience connectivity

Stimulants are one of the oldest, most potent, and most broadly used prescription psychoactive drugs with 14 million users^1,4,5^ and over $2.2 billion annual sales in the United States,^143^ but their effects on the brain remain incompletely understood with divergent prior findings.^42^ Recent advances including large brain-wide association study (BWAS) datasets^90,119^ and precision imaging drug trials (PIDT) for controlled verification of BWAS findings^86,88,123^ allowed us to investigate the brain effects of stimulants on a scale not previously possible. Capitalizing on the recognition of action regions embedded in primary motor cortex,^109^ comparison with drug target receptor maps,^63,138^ and data-driven statistical approaches^121,122^ allowed us to resolve previously ambiguous findings. This multi-modal approach revealed that the largest stimulant-related changes in functional connectivity (FC) are in action/motor regions reflecting arousal state,^112–114^ and in tightly-coupled salience and parietal memory networks (SAL/PMN) associated with anticipation of reward/aversion and action-relevant memory.^79–81^

### Stimulants have little direct effect on attention

Prior theories regarding prescription stimulants posited direct beneficial effects on attention and control networks intersecting prefrontal cortex such as DAN, VAN, and FPN.^21^ There is evidence that prefrontal cortex is associated with attention deficit in ADHD^144^ and modulated by catecholamines.^31,32^ However, much prior neuroimaging evidence that stimulants act primarily on attention and control networks comes from studies using region of interest (ROI) methodologies focused on these a priori networks.^43,44,46–48,64^ Increased computational power and advances in statistical modeling now enable comparison of the relative effects of stimulants on different networks,^96–98^ see our Supplemental Discussion. Here, with a large sample of children (*n* = 5,795), we found no significant differences in DAN, VAN, or FPN related to stimulants after accounting for larger stimulant-related differences in other brain networks (see Supplemental Table 2 for a power analysis). We found correspondingly no significant difference in performance on the NIH Toolbox or n-back, tasks involving attention and working memory, in healthy children taking stimulants. Instead, performance of children with ADHD taking a stimulant improved to the level of the rest of the cohort. These imaging and behavioral evidence do not support the hypothesis that the primary effect of stimulants is to increase attentional ability through direct modulation of attention and control networks.

Instead, the largest stimulant-related differences in cortical FC were in somato-cognitive action and motor regions. Attempting to reconcile stimulant effects in motor cortex with their use in treating ADHD, it has been argued that stimulants might reduce motoric output by enhancing cortical inhibition in motor cortex.^82,103–108^ While inhibition of motoric output might be desirable when stimulants are taken to treat ADHD, stimulants are also effective in contexts where the goal is to increase motoric output, such as athletic enhancement.^19^ We observed that stimulant-related differences in motor cortex FC were highly concordant with the FC pattern of getting more sleep or being more alert. Thus, the role of stimulants in action and motor cortex could be related to increased sympathetic drive and higher arousal, consistent with recent insights into action and motor cortex function.^86,112–118^

The seemingly paradoxical effect that stimulants can reduce hyperactivity may instead be related to their dopaminergic effects on salience processing. The second largest stimulant-related differences in FC were in SAL/PMN which, together, are thought to encode anticipated reward/aversion and thus influence the decision to persist at a task or switch to a more rewarding task.^22,53,68,69,71,72,83,145–150^ Aspects of ADHD hyperactivity could be associated with searching for more rewarding actions and thus better undestood as motivational than motoric. We hypothesize that stimulants reduce task-switching and thus appear outwardly to facilitate attention by elevating the perceived salience of mundane tasks (e.g. math homework)^35^ through their effect on SAL, with a complementary process boosting memory through PMN. Both BWAS and controlled PFM data reported here are consistent with prior behavioral studies describing the influence of prescription stimulants on persistence^27,28^ and effort,^30,33,78^ compensating for unfocused attention in individuals with ADHD without affecting cognitive ability.^27,28,33,34^ Although beyond the scope of this study, future work should assess whether stimulants increase task-fMRI activation in response to smaller anticipated rewards.

### Stimulants rescue brain connectivity from short-term sleep deprivation

Stimulants increase synaptic norepinephrine,^2,3^ promoting arousal and wakefulness.^8,9,20,151,152^ We observed stimulant-related differences in sensorimotor FC aligned with norepinephrine receptor density, consistent with recent insights into action and motor cortex function.^86,112–118^ Remarkably, we found that taking a stimulant before scanning made the brain connectivity of children with less sleep indistinguishable from that of well-rested children. Stimulants also rescued cognitive performance in children with less sleep. Thus, stimulants appeared to rescue the brain from the effects of sleep deprivation, at least in the short term. The ability of stimulants to rescue cognitive decrements in sleep-deprived individuals through modulation of the brain’s arousal system may be an important reason why many purported cognitive advantages of stimulants do not replicate in controlled experimental cohorts with little variation in sleep.^27,28,33,34,152^

While our results appear to show that the cognitive performance of sleep-deprived children benefited from stimulants, we caution that mounting evidence points to cumulative health consequences of long-term sleep deprivation including increased risk of depression, cellular stress, and neuronal loss.^101,153^ A wash-out study collecting fMRI data in sleep-deprived participants shortly after taking stimulants and later after drug levels have fallen could assess whether the beneficial effects of stimulants persist or reverse after drug concentrations taper off in the afternoon. Additional long-term studies are needed to evaluate whether stimulant users are less likely to get adequate sleep and measure the cumulative effects of sleep loss over the lifespan.

### Patients with ADHD benefit from stimulants

ADHD is the primary medical indication for stimulants.^6^ ADHD is a heterogeneous condition with reported changes in attention networks, salience networks, mixed mechanisms,^21,22,53,68,69,154,155^ and even the existence of distinct ADHD subtypes,^156–158^ including evidence from the ABCD Study.^154,155^ Our findings show that stimulants improve school grades and cognitive performance in children with ADHD without increasing cognitive ability or bestowing any unfair advantage.^17^ We also show that FC differences related to stimulants are similar to those of getting more sleep, and that getting more sleep was itself associated with increased cognitive performance.^102^ Sleep disturbance is a common comorbidity of ADHD and a common complication of stimulant treatment,^159^ therefore clinicians should screen for sleep disturbance in children with ADHD both before and after prescribing a stimulant.

### Stimulants increase drive, not attention

Understanding which brain systems are affected by stimulants is important both to guide treatment decisions and facilitate development of novel psychoactive drugs. Using resting-state fMRI, we showed that stimulants mimic the effects of sleep (arousal) and reward expectation (salience) through motor/arousal and salience/memory networks consistent with boosting effort^30,33,71,78^ and persistence,^27,28^ not attentional nor cognitive ability. The beneficial effects of stimulants on motivation and persistence are consistent with their many uses beyond the treatment of ADHD including to treat narcolepsy,^7^ promote wakefulness after traumatic brain injury,^8,9^ increase diet adherence,^11–13^ and enhance athletic performance.^19^ Some of the benefits of stimulants could also be attained by getting sufficient sleep each night,^152^ something about half of children^100,101^ and adults^160^ go without. Any additional stimulant-specific effects not shared with being better rested may derive from elevating the perceived salience of goal-directed actions (SAL) and memories (PMN). Thus, stimulants seem to boost our ability to persist in drudgery, without significantly affecting intrinsically rewarding tasks.

## Methods

### Ethics

This project used resting-state functional MRI, demographic, biophysical, and behavioral data from 11,572 8–11 year old participants from the ABCD 2.0 release.^120^ The ABCD Study obtained centralized institutional review board (IRB) approval from the University of California, San Diego. Each of the 21 sites also obtained local IRB approval. Ethical regulations were followed during data collection and analysis. Parents or caregivers provided written informed consent, and children gave written assent. Data from the Psilocybin PFM study^123^ were collected in accordance with protocols approved by the Washington University in St. Louis IRB. This project also includes published derivatives from other studies^113,117,118,137^ whose protocols were governed by their respective IRBs.

### Behavioral

The Adolescent Brain Cognitive Development (ABCD) study participants are well-phenotyped with demographic, physical, cognitive^161^, and mental health^162^ batteries. We used the NIH Toolbox^141^ and parent reported school grades as measures of out-of-scanner cognitive ability. Data were downloaded from the NIMH Data Archive (ABCD Release 2.0), and the traits of interest were extracted using the ABCDE software we have developed and which we have made available here: https://gitlab.com/DosenbachGreene/abcde.

### Prescription Stimulant Medications

The ABCD Study asked parents to recall their children’s prescription medications. Parents searched for their children’s medications on an interactive tablet linked to the RxNorm database.^163^ Parents were also asked whether their child took the medication in the last 24 hours. Stimulants are dosed in the morning, therefore children whose parents reported giving stimulants within the last 24 hours were assumed to have taken the stimulant on the morning of their MRI scans. Complete information about dosage and formulation (e.g. tablet, liquid, extended release) were not available for the first year of the study. Using the ABCDE software, we cross-referenced parent responses with the RxNorm database to identify children taking a drug with one of the following active ingredients: methylphenidate, dexmethylphenidate, amphetamine, dextroamphetamine, or lisdexamfetamine. The stimulant drug serdexmethylphenidate was approved by the FDA in 2021, after the first year of ABCD data had been collected. Among the sample of 5,795 children with complete data, 7.1% (73.6% boys) were prescribed a stimulant and 5.8% (73.0% boys) took the stimulant on the day of scanning (*n* = 337).

### ADHD

Several algorithms have been proposed to identify children with ADHD in the ABCD Study.^124^ This study used the stringent “Tier 4” criteria from Cordova et al.^124^ These criteria include children who met criteria for ADHD “present” or “current” on the Kiddie Schedule for Affective Disorders and Schizophrenia (KSADS-COMP).^164^ Children with intellectual disability, bipolar disorder, schizophrenia or psychotic symptoms were excluded. Children who scored below clinical cutoff on the teacher-reported Brief Problem Monitor (BPM) scale,^165^ or who scored below clinical cutoff on the parent-reported Child Behavioral Checklist (CBCL) attention or ADHD scales^166^ were also excluded. Children with missing data were not excluded.

Among the sample of 5,795 children with complete data, 3.0% of children (66.9% boys) had ADHD of whom 43.4% were prescribed a stimulant and 34.9% took a stimulant on the day of scanning. Conversely, 18.1% of children taking a stimulant on the day of scanning had ADHD. To reconcile this paradox, we defined a less stringent criteria for ADHD used for exploratory analysis only; the stringent criteria was used for our main analyses. The less stringent criteria was based on the KSADS-COMP only and included children with ADHD “present,” “past,” “in remission,” or of an “unspecified” subtype. A majority (75.0%) of children taking stimulants met these less stringent, exploratory criteria for ADHD.

### Sleep

Parents were asked questions about their child’s sleep disturbances.^134^ We reversed the order of the responses to create a monotonically increasing scale of average sleep duration with 1 = less than 5 hours, 2 = 5-7 hours, 3 = 7-8 hours, 4 = 8-9 hours, and 5 = greater than 9 hours. We used average sleep duration as a surrogate measure of arousal/wakefulness at the time of scanning.

### Covariates

Following published guidelines for the ABCD Study,^167,168^ we selected average in-scanner head motion (FD), age (in months), sex (assigned at birth), household income bracket, highest level of education achieved by a parent, and whether or not parents were married as nuisance covariates. The marriage covariate was supplemented by an additional covariate describing whether there was one or more than one adult caregiver in the household (regardless of marital status). There is controversy regarding inclusion of race or genetic ancestry as a default covariate;^168^ we did not include race as there is no biologically plausible mechanism by which it would affect the brain’s response to stimulant medications.

We selected additional covariates relevant to our hypotheses, including ADHD diagnosis (using the stringent “Tier 4” criteria above).^124^ Diurnal variations are reported to affect FC,^169^ therefore we also included time of scan (morning or afternoon), and day of week (weekday or weekend). Except where otherwise noted, sleep duration was included as a covariate in analyses of stimulants, and stimulant taking was included as a covariate in analyses of sleep duration.

### MR imaging

Functional magnetic resonance imaging (fMRI) was acquired at 21 sites using a protocol harmonized for 3 Tesla GE, Philips, and Siemens scanners with multi-channel receive coils.^128^ In addition to anatomical and task-fMRI, each participant had up to four 5-minute-long resting-state scans (TR = 800 ms, 20 minutes total). A subset of sites using Siemens scanners used FIRMM motion tracking software that allows extending the scan on the basis of on-line measurement of motion.^170^

Following acquisition, fMRI data were processed using standardized methods including correction for field distortion, frame-by-frame motion co-registration, alignment to standard stereo-tactic space, and extraction of the cortical ribbon.^171^ Resting-state data were further processed to remove respiratory and motion artifact by temporal bandpass filtering, global signal regression, and regression against the rigid-body motion parameters using the ABCD-BIDS motion processing pipeline,^172,173^ a derivative of the Human Connectome Project (HCP) processing pipeline.^174^ Processing dependencies included FSL^175^ and FreeSurfer.^176^ Functional MRI data acquired at different study sites were harmonized using CovBat.^177–179^

### Parcellation

It is possible to compute functional connectivity between each voxel or vertex. However, this approach is burdened by a high proportion of unstructured noise and large computer memory requirements. We therefore adopted a parcel-based approach based on the 333 cortical parcels described by Gordon and Laumann^133^ augmented by the 61 subcortical spheres described by Seitzman^180^ for a total of 394 parcels, or nodes.

### Removing head motion artifact

Motion in fMRI studies is typically estimated using spatial co-registration of each fMRI volume (or frame) to a reference frame.^181^ In this study we quantified motion using framewise displacement, FD (L1-norm), in millimeters, after filtering for respiratory artifact.^172,182^ Exclusion of frames with FD > 0.2 mm has been shown to reduce spurious findings associated with residual motion artifact in high-motion groups,^183,184^ such as children with ADHD. Participants with less than 8 minutes (600 frames) of resting-state data remaining after motion censoring, the minimum duration needed for high-quality estimation of connectivity,^85^ were excluded from analysis. Of the 11,875 children recruited in the first wave of the ABCD Study, 8,486 had more than 8 minutes of rs-fMRI data after censoring frames with FD > 0.2 mm.

### Motion impact assessment

After removing head motion artifact, we quantified the impact of residual head motion artifact on our brain-behavior associations of interest, stimulant taking, and sleep duration, using the SHAMAN method.^184^ The covariates described above were included as regressors of non-interest.

### Functional connectivity

We employed standard approaches for computing functional connectivity. The methods are briefly summarized here. By convention, each brain region or parcel is referred to as a node. The functional connections between nodes, which are referred to as edges, are computed as the pairwise linear correlation coefficients between nodes. As correlations are constrained to vary from -1 to 1, the correlation coefficients were Fisher Z transformed (inverse hyperbolic tangent function) to lie on an approximately normal distribution. Ordinary least squares (OLS) regression was performed independently at each edge.

### Marginal model and bootstrapping

The ABCD data are clustered by study site and family (some participants are siblings). In addition to data harmonization across sites with CovBat,^177–179^ we explicitly modeled site differences and sibling relationships in our statistical analyses. A linear mixed-effects model with site and family random effects would have been computationally expensive due to the large number of participants and features (edges) in this study. Instead, we computed edgewise cluster-robust marginal *t* -values corrected for site and family using the sandwich estimator^185,186^ followed by wild bootstrap under the null model with the Rademacher distribution.^187^ This approach has been shown to yield comparable results to mixed effects regression at lower computational cost in large neuroimaging datasets.^173,188^

### Network level analysis

We are principally interested in FC differences involving canonical networks (e.g. DMN, VIS, etc.), not differences involving specific edges. Network Level Analysis (NLA) is an adaptation of enrichment analysis that performs inference at the level of canonical networks. We performed NLA using previously described methods.^121,122^ Briefly, FC values at each edge were studentized using the cluster-robust sandwich estimator approach described above^185,186^ to obtain edge-level FC *t* -values. The average FC *t* -value of edges within each network pair (e.g. DMN and VIS) was compared with the average FC *t* -value value over the whole connectome using Welch’s *t* -test.^189^ A Welch’s *t* -value of zero indicated no difference in FC relative to the whole connectome. A positive Welch’s *t* -value indicated enrichment of FC differences within a network pair, i.e. a large change in connectivity. A negative Welch’s *t* -value indicated depletion of FC differences within a network pair, i.e. a small change in connectivity.

Separately, whole networks were compared to the connectome by averaging the absolute values of the FC *t* -values in each network. Positive Welch’s *t* -values indicated enrichment of FC differences within a network, i.e. a large change in connectivity.

Inference was performed by generating a null distribution of Welch’s *t* -values using the same wild bootstrap procedure described above^187^ with 2,000 bootstrap iterations. The Westfall-Young step-down procedure^127^ was used to control the family wise error rate (FWER) from comparisons across multiple network pairs.

NLA is biased toward detection of significant changes in large networks pairs with many edges. Therefore, related networks with a small number of nodes were combined as indicated in Figure 1 for the purpose of statistical inference. Lateral and medial visual networks were combined into a single visual network. Salience and parietal memory networks were combined into a single SAL/PMN network. Premotor, somatomotor hand, somatomotor mouth, somatomotor foot, and somatocognitive action networks were combined into a single SM network.

### Power analyses

Like all methods to account for multiple statistical comparisons, network level analysis (NLA) controls the false positive error rate at the expense of false negative error rate, or statistical power. A prior study^45^ on stimulant-related FC differences within attention networks with *n* = 24 participants reported a *t* -value of 4.35 corresponding to an effect size (Cohen’s *d* ) of 0.89. We assessed the power of our NLA approach to detect an FC difference of this size within attention or control networks: DAN, VAN, or FPN. For each network, we simulated a Welch’s *t* -value using the formula:

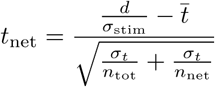

Where:

- *t*_net_ is the Welch’s *t*-statistic for a network
- *d* is the effect size, e.g. 0.89
- *σ*_stim_ = 0.058 is the standard error of regression for stimulants, i.e. the square root of the diagonal element in (*X*^⊤^*X*)^−1^
- *̄t* = 0.031 is the average *t* -statistic for all edges in the connectome
- *σ_t_* = 1.35 is the standard deviation of *t* -statistics in the connectome
- *n*_tot_ = 77,421 is the total number of edges in the connectome
- *n*_net_ is the number of within-network edges (DAN: 496, VAN: 253, FPN: 276)

The simulated Welch’s *t* -value, *t*_net_, was ranked against the bootstrapped null distribution of Welch’s *t* -values to compute a *P* -value. The *P* -value was corrected for multiple comparisons using the Westfall-Young step-down procedure. Power to detect an effect size *d* within the given network was calculated as 1 - *P*. See Supplemental Table 2 for minimum detectable effect sizes at different power levels.

### Generation of brain maps

Analysis of rs-fMRI data was performed on (394^2^-394)/2 = 77,421 distinct edges arising from the 333 Gordon-Laumann cortical parcels and 61 Seitzmann subcortical spheres.^133,180^ Some results were projected back into the space of the 333 cortical parcels for visualization as brain maps. Seed-based FC maps were generated from an exemplar seed parcel in somatomotor hand region, which was selected *a posteriori* as the parcel with the greatest difference in FC related to stimulants. Brain maps of magnitude difference in FC were generated by computing the root mean square (RMS) average change in FC for each row in the FC matrix. The RMS values were rendered on their corresponding cortical parcels using Connectome Workbench.^190^

In Supplemental Figure 3 the vertex-wise nucleus accumbens seed map was generated using the subcortical volume for nucleus accumbens from the Human Connectome Project^135,174^. In Supplemental Figure 17, the value at each cortical parcel was computed as the linear (Pearson) correlation between each row in the stimulant FC matrix with each corresponding row in the sleep FC matrix.

### Statistical Comparison of Brain Maps

Inference on similarity between brain maps was performed using the rotational null model of Vázquez-Rodríguez for parcellated surface maps^129,191^ using the NeuroMaps software.^130^ Comparisons were performed for the 333 parcels on the cortical surface^133^ only. Many maps were thresholded at 50% intensity for visual presentation (e.g. Figure 4), but the whole range of intensity values were used for quantifying similarity. First we calculated the real correlation *r* between two maps across the 333 parcels. Then we generated 2,000 rotational null maps and computed the correlations *r* _Ø,1_,… *r* _Ø,2000_ between each pair of null maps. Finally, we computed the p-value for the two-tailed alternative hypothesis of *r* ≠ 0 by counting the number of permutations in which |*r* | < |*r* _Ø_| and dividing by the total number of permutations.

### Task-fMRI Analysis

The n-back task was used in the ABCD Study to engage working memory and cognitive control in adolescents.^128^ There was less fMRI data available for the n-back task compared to rest due to greater scan time allocated to resting-state data acquisition; consequently, there were only *n* = 1,944 children with high-quality n-back data (FD < 0.2 mm and greater than 8 minuts of scan time) of whom *n* = 109 took a stimulant on the day of scanning. N-back data were analyzed in two ways. First, data were treated as rest, without regressing out the task paradigm, to test the hypothesis that stimulants would affect FC during an attention-demanding task differnetly than they would at rest, see Supplemental Figure 4. Second, we performed conventional task-fMRI analysis for the 0-back vs fixation contrast using FSL’s FEAT with default settings,^175,192,193^ see Supplemental Figure 5.

### Replication in healthy adults

Five healthy adults without ADHD ages 18-45 years (2 male, 3 female) participated in a randomized cross-over pharmacometric fMRI study in which participants received methylphenidate 40 mg or psilocybin 25 mg on separate days in a random order.^123^ (A sixth participant taking a prescription stimulant was excluded from analysis.) Image acquisition was divided across multiple days. Resting-state fMRI was acquired using the protocol below with multiple 15-minute-long rs-fMRI scans per day of scanning. Each participant underwent at least 4 baseline scans before receiving either methylphenidate or psilocybin.

The fMRI acquisition protocol was similar to ABCD. We used an echo-planar imaging sequence with 2 mm isotropic voxels, multi-band 6, multi-echo 5 (TEs: 14.20 ms, 38.93 ms, 63.66 ms, 88.39 ms, 113.12 ms), TR 1761 ms, flip angle = 68°, and in-plane acceleration (IPAT/grappa) = 2. This sequence acquired 72 axial slices (144 mm coverage). Each resting scan included 510 frames (lasting 15:49 minutes) as well as 3 frames at the end used to estimate electronic noise. Data were processed using a previously-described custom pipeline^123^ except that we performed global signal regression to more closely match the ABCD data.

Data were co-registered to the same atlas as the ABCD data and parcellated using the same 394 parcellation used for the ABCD data. A 394 × 394 FC matrix was computed for each rs-fMRI scan using the methods above and motion censoring threshold of FD < 0.2 mm as in the ABCD data. An edge-wise linear mixed effects model was used to compare scans on methylphenidate to baseline scans. The data on psilocybin were not used. Sex was modeled as a fixed effect. The model included a random intercept for scan session (a day of scanning) as well as a random intercept and slope (for methylphenidate) within participant. Due to the small number of participants (*n* = 5), we did not attempt to perform permutation-based significance testing or network level analysis. Edge-wise *t* -values are reported in Figure 11 and were used to generate the cortical surface maps shown in Figure 2.

### Norepinephrine transporter data

PET maps were compiled by Hansen et al.^138^ and projected onto the cortical surface using Connectome Workbench.^190^ The map of norepinephrine transporter was generated using the 11C-MRB (methylreboxetine) ligand (*n* = 20).^137^ The supplemental dopamine receptor map for D1 was generated using the 11C-SCH23390 ligand (*n* = 13).^194^ The D2 map was generated using the 11C-FLB457 ligand (*n* = 6).^195^

The data were downloaded from: https://github.com/netneurolab/hansen_receptors

### Independent arousal data

The ABCD Study does not include physiologic arousal data, therefore we compared FC differences related to sleep duration (a surrogate measure of arousal) in ABCD data to physiologic arousal maps from two independent studies, see Figure 4.

The EEG alpha slow wave index arousal template (*n* = 10)^117,118^ was projected onto the cortical surface using Connectome Workbench.^190^

The data were downloaded from: https://github.com/neurdylab/fMRIAlertnessDetection

The respiratory variation arousal map (see Figure S2B “PLV_magnitude” from Raut et al.)^113^ was generated from *n* = 190 participants with simultaneous fMRI and respiratory (chest bellows) data in the WU-Minn Human Connectome Project (HCP) 1200 Subject Release.^135^

The data were downloaded from: https://github.com/ryraut/arousal-waves

The arousal map is found under: output_files/HCP_RV_coherencemap.dtseries.nii

## Data availability

Participant level data from ABCD are openly available pursuant to consortium-level data access rules. The ABCD data repository grows and changes over time (https://nda.nih.gov/abcd). The ABCD data used in this study came from ABCD Annual Release 2.0 (https://doi.org/10.15154/1503209).^120^

### Code availability

Code for Network Level analysis can be found at: https://github.com/WheelockLab/NetworkLevelAnalysisBeta
Code for processing ABCD and UKB data can be found at: https://github.com/DCAN-Labs/abcd-hcp-pipeline

## Acknowledgements

ABCD acknowledgement

Data used in the preparation of this article were obtained from the Adolescent Brain Cognitive Development (ABCD) Study (https://abcdstudy.org), held in the NIMH Data Archive (NDA). This is a multisite, longitudinal study designed to recruit more than 10,000 children age 9-10 and follow them over 10 years into early adulthood. The ABCD Study is supported by the National Institutes of Health and additional federal partners under award numbers U01DA041022, U01DA041028, U01DA041048, U01DA041089, U01DA041106, U01DA041117, U01DA041120, U01DA041134, U01DA041148, U01DA041156, U01DA041174, U24DA041123, U24DA041147, U01DA041093, and U01DA041025. A full list of supporters is available at https://abcdstudy.org/federal-partners.html. A listing of participating sites and a complete listing of the study investigators can be found at: https://abcdstudy.org/scientists/ workgroups/. ABCD consortium investigators designed and implemented the study and/or provided data but did not necessarily participate in analysis or writing of this report. This manuscript reflects the views of the authors and may not reflect the opinions or views of the NIH or ABCD consortium investigators.

The ABCD data repository grows and changes over time. The ABCD data used in this report came from Anual Release 2.0, DOI 10.15154/1503209.^120^

## Precision imaging drug trial data

Data for the precision imaging drug trial validation of methylphenidate in 5 healthy adults was collected by Joshua Siegel and others as part of a larger study on the brain effects of psilocybin.^123^ The data are available at: https://wustl.box.com/v/PsilocybinPFM

## EEG/fMRI arousal template data

Data for the EEG/fMRI arousal template were collected and made publicly available by Catie Chang and others in the Neuroimaging & Brain Dynamics Lab at Vanderbilt University.^117,118^ The data were downloaded from: https://github.com/neurdylab/fMRIAlertnessDetection

## Respiratory variation arousal data

Data for the respiratory variation arousal map were collected at the University of Minnesota and Washington Unviersity in St. Louis as part of the Human Connectome Project (HCP) 1200 Subject Release.^135^ The respiratory variation arousal map was generated by Ryan Raut and others at WU.^113^ The data were downloaded from: https://github.com/ryraut/arousal-waves

## Positron emission tomography data

Data for positron emission tomography (PET) maps of receptor density were compiled by Justine Hansen and others at the Montréal Neurological Institute. These data include norepinephrine,^137^ D1,^194^ and D2^195^ receptor densities. The data were downloaded from: https://github.com/netneurolab/hansen_receptors/

## Supplemental Figures & Tables

**Figure 1:**
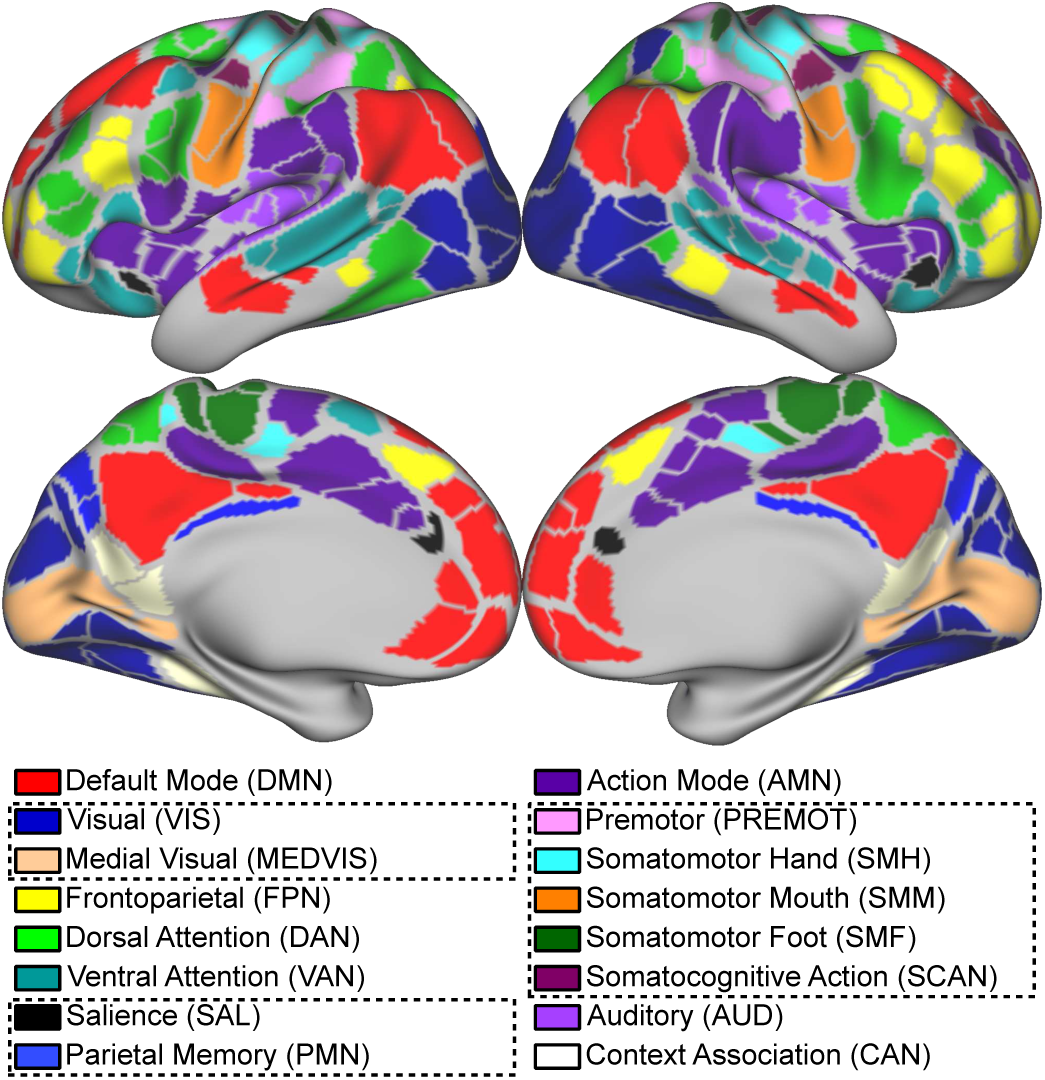
Network communities shown on the Gordon-Laumann 333 cortical parcels.^133^ Networks grouped together for network level analysis are indicated by boxes.

**Figure 2:**
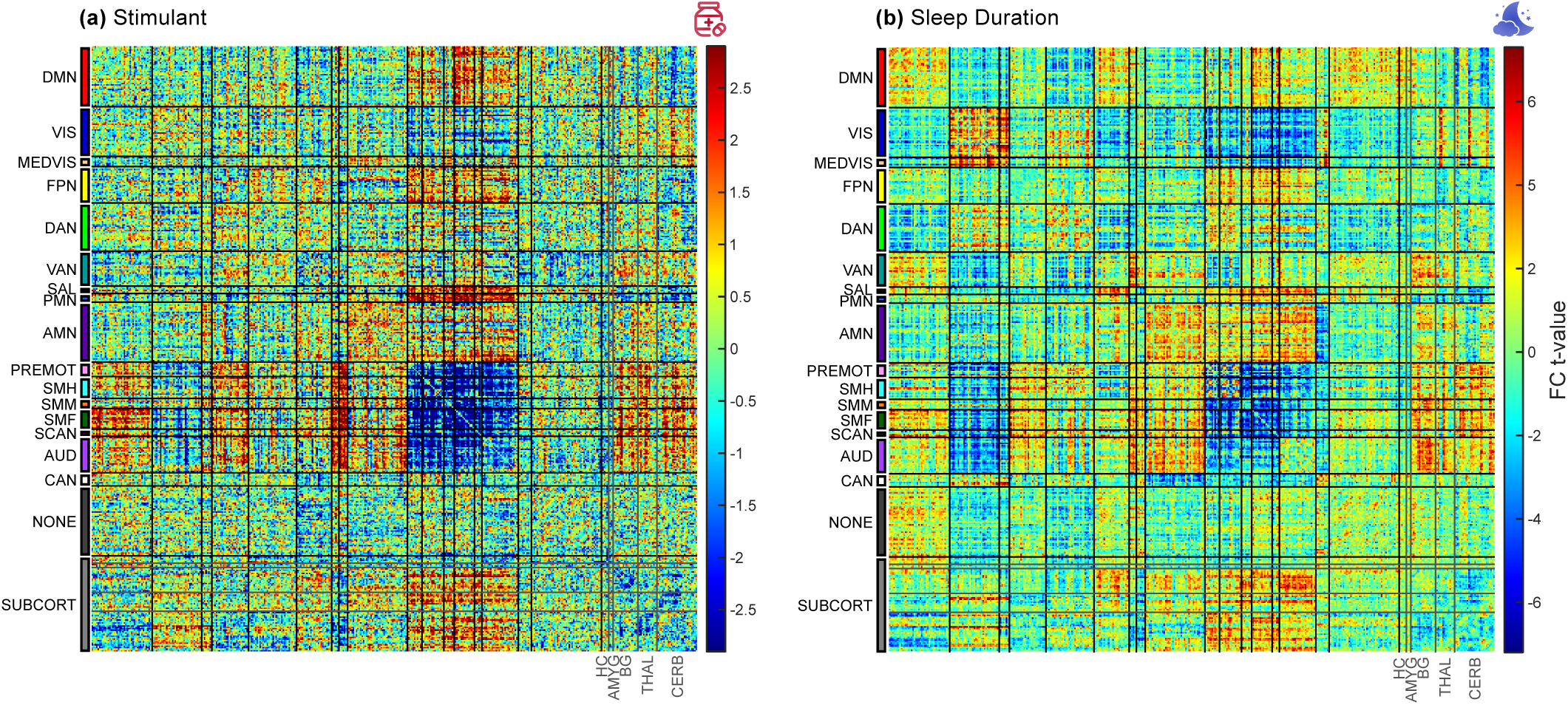
Differences in FC related to stimulants and sleep. **(a)** Differences in FC related to taking a stimulant on the day of scanning (*n* = 5,795, 337 taking stimulants). **(b)** Differences in FC related to sleep duration (*n* = 5,795). Effect size and *t* -values were overall greater for sleep duration. The FC matrices are edge-for-edge correlated at *r* = 0.38. For names and locations of networks see Supplemental Figure 1.

**Figure 3:**
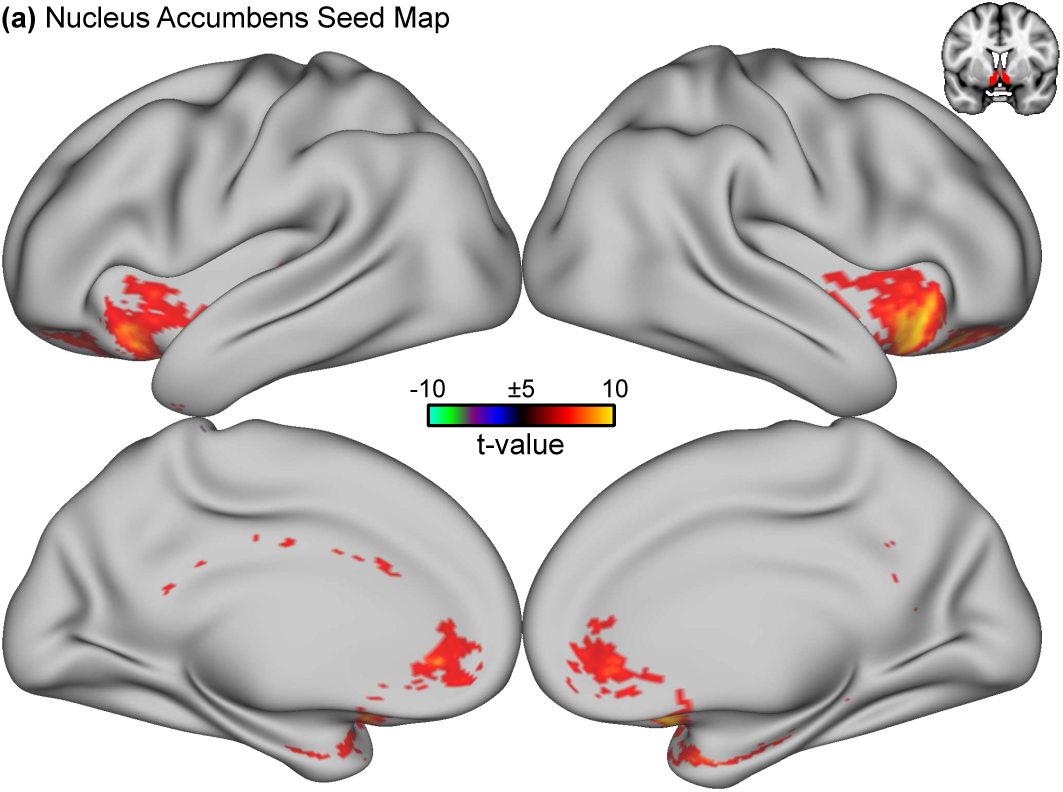
Nucleus accumbens seed map. FC (t-value) between nucleus accumbens and cortex is shown for all *n* = 5,795 participants in ABCD. Accumbens FC overlaps with right anterior inferior insula, the hub of the salience network. The nucleus accumbens seed is shown at top right. There was no significant difference in accumbens-cortex FC related to taking stimulants.

**Figure 4:**
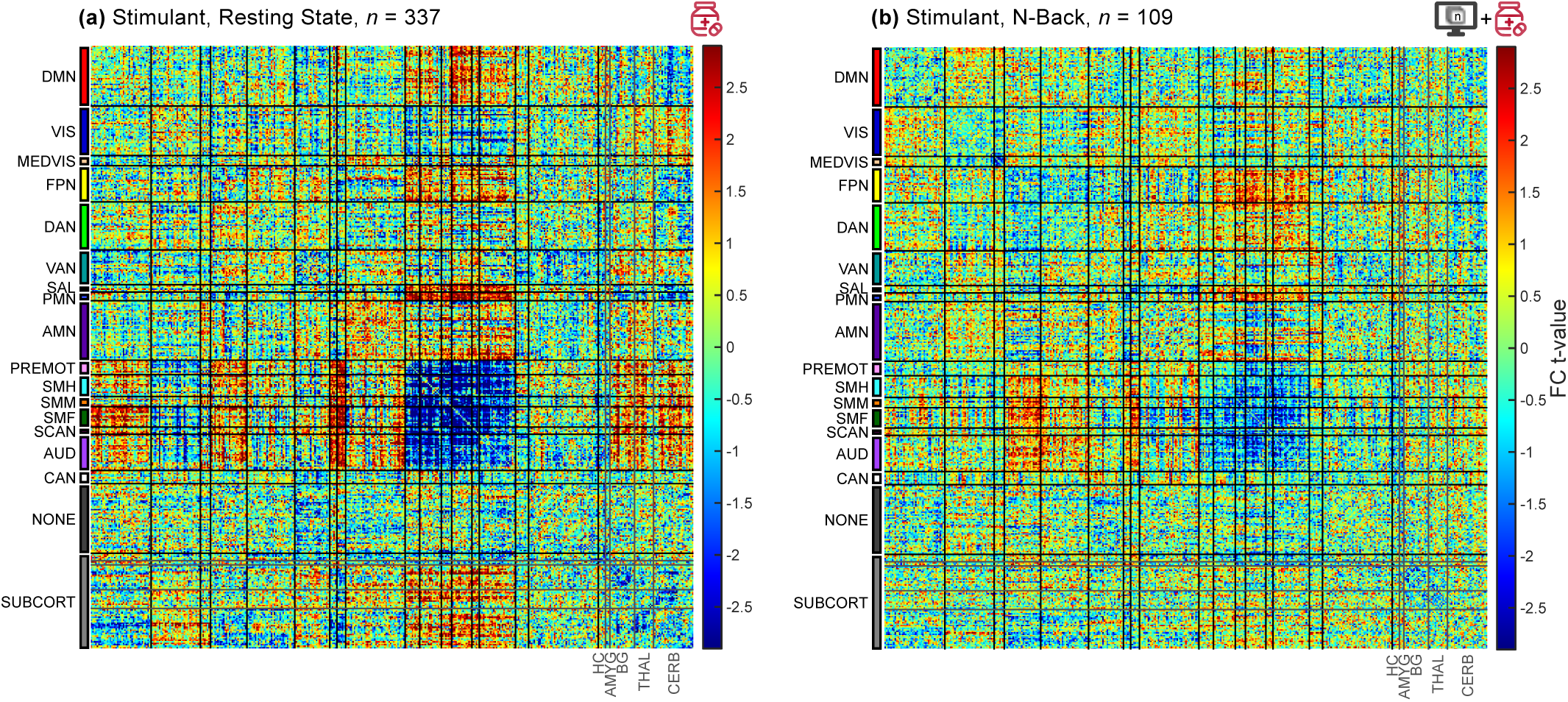
Differences in FC related to stimulants in resting and task (n-back) data. **(a)** Stimulant-related differences in FC during resting state with *n* = 337 participants taking stimulants among a total of *n* = 5,795 participants, as in Supplemental Figure 2. **(b)** Stimulant-related differences in data collected during the n-back task treated as rest with *n* = 109 children taking stimulants and *n* = 1,944 children total. The task paradigm was not regressed out. The FC matrices are edge-for-edge correlated at *r* = 0.26. For names and locations of networks see Supplemental Figure 1.

**Figure 5:**
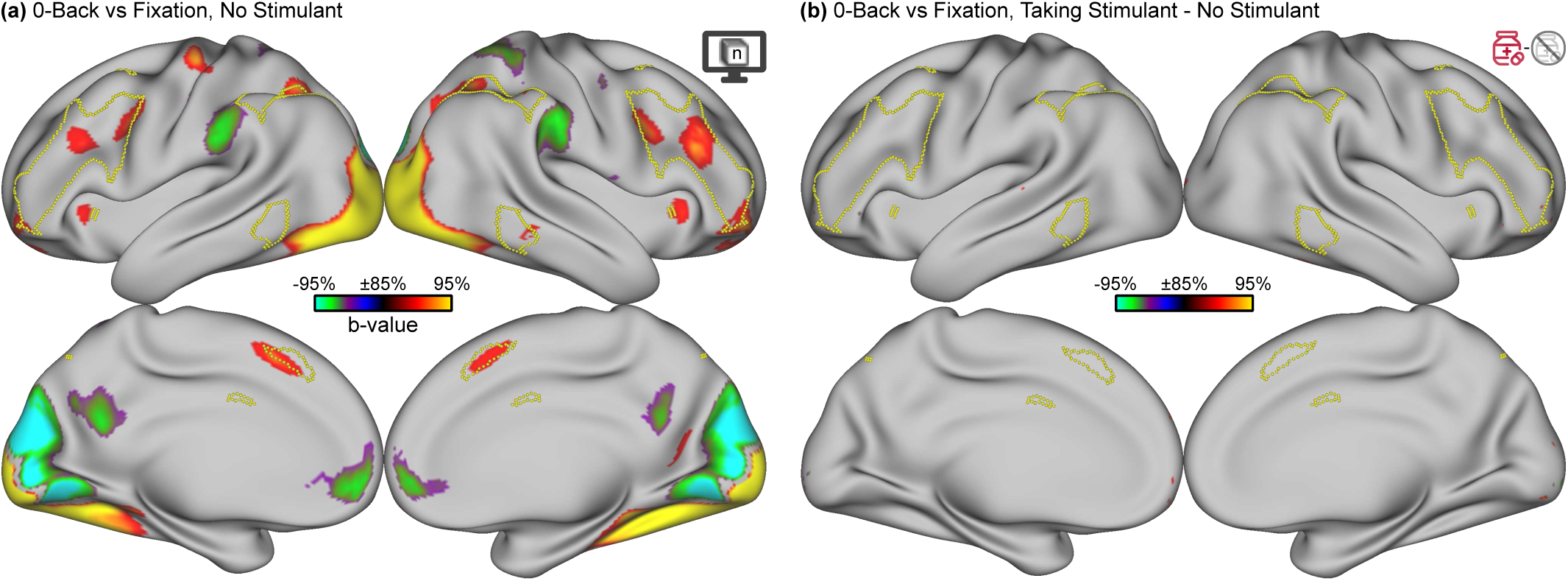
Task-evoked activation for the n-back. **(a)** Task-evoked activation for 0-back vs fixation contrast in all participants. Regression coefficients (b-values) between 85% and 95% of the maximum are shown. The frontoparietal network (FPN) is outlined in yellow. There was less fMRI data available for the n-back task compared to rest due to greater scan time allocated to resting-state data acquisition; consequently, there were only *n* = 1,944 children with high-quality n-back data. **(b)** Higher-order contrast for 0-back vs fixation in children taking stimulants (*n* = 109) vs not taking stimulants, shown on the same b-value scale.

**Figure 6:**
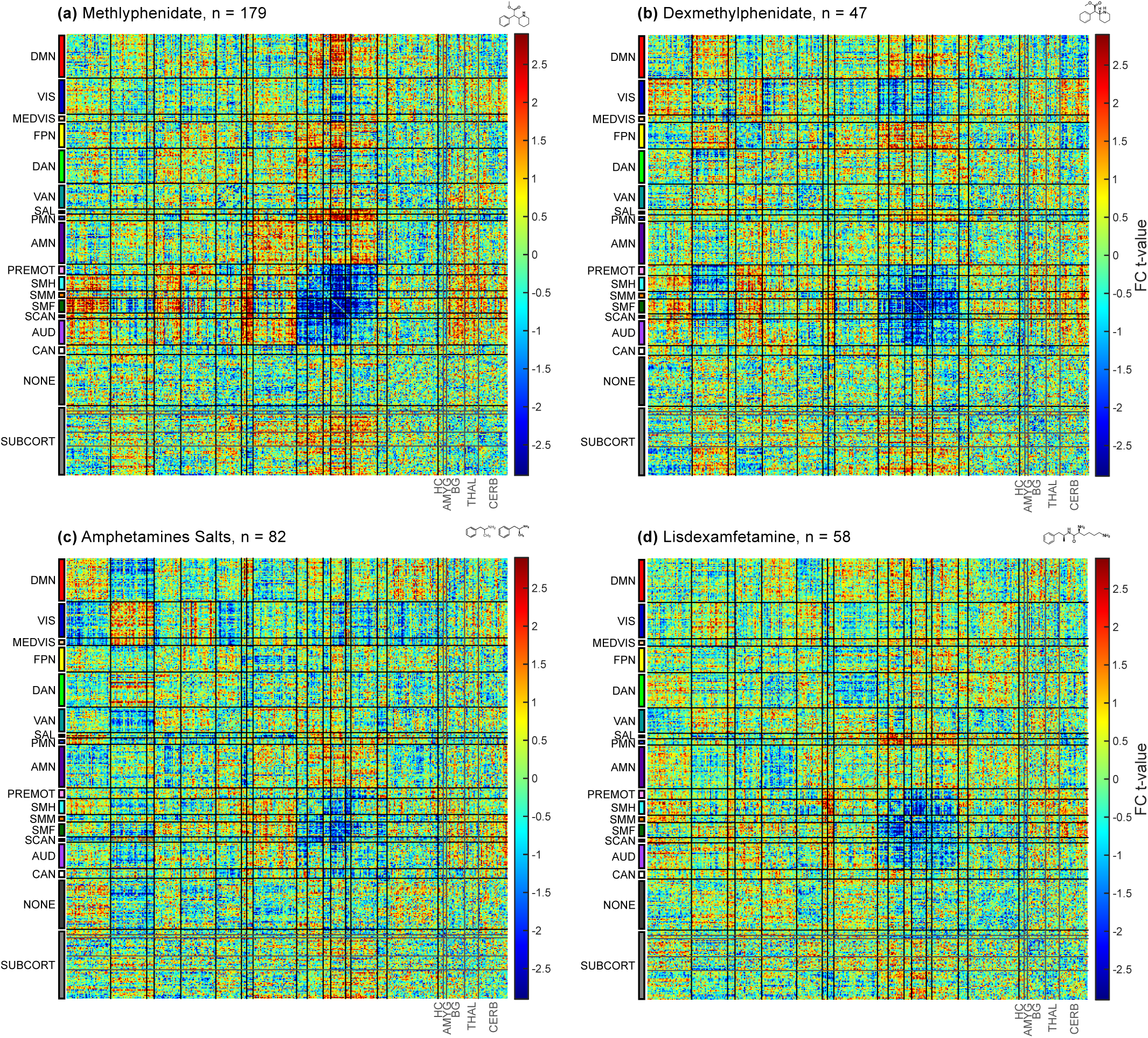
Differences in FC related to different stimulant drugs. **(a)** Methylphenidate (Ritalin), *n* = 179 children. The FC matrix for methylphenidate was edge-for-edge correlated with the pooled FC matrix for all stimulants at *r* = 0.81. **(b)** Dexmethylphenidate (Focalin), *n* = 47, *r* = 0.53. **(c)** Mixed amphetamine salts (Adderall), *n* = 82, *r* = 0.54. **(d)** Lisdexamfetamine (Vyvanse), *n* = 58, *r* = 0.48. For names and locations of networks see Supplemental Figure 1.

**Figure 7:**
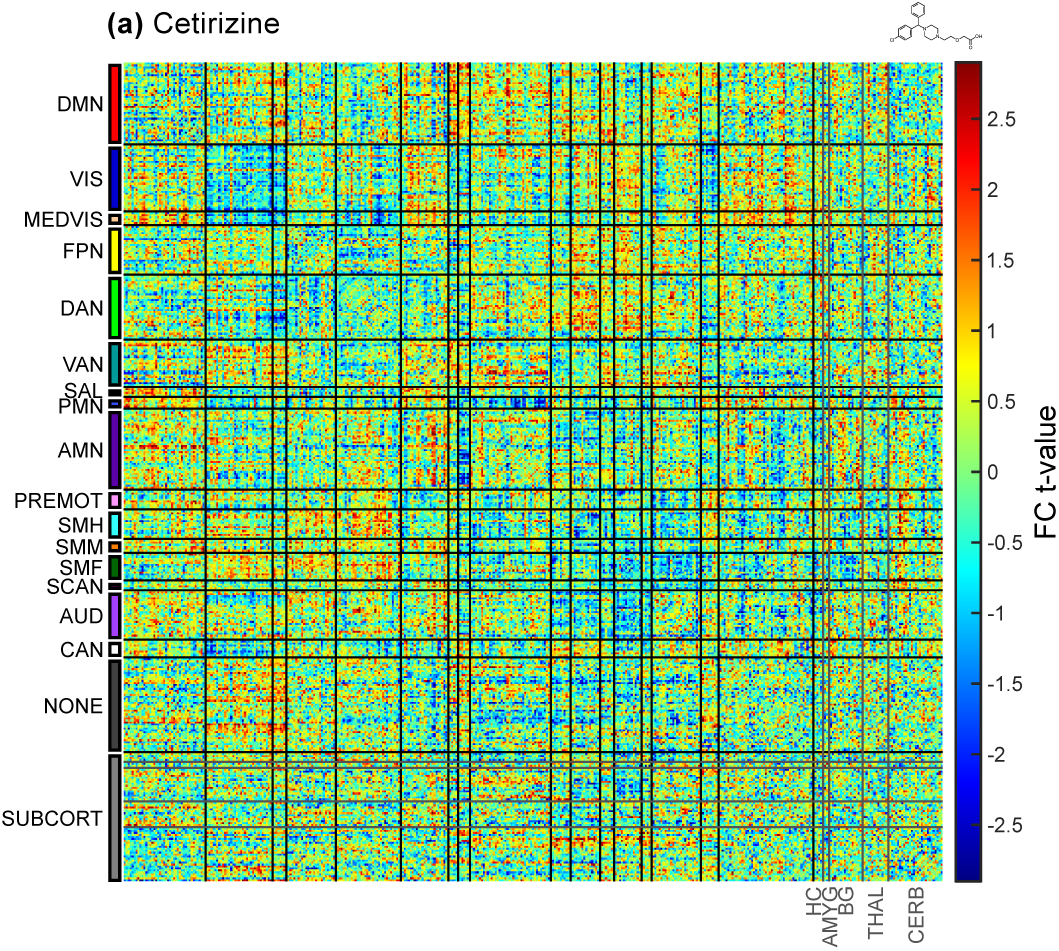
Differences in FC related to cetirizine. Cetirizine, a commonly-taken allergy medication without psychoactive properties,^132^ was selected as a negative control. FC differences for children taking cetirizine within 24 hours of scanning are shown with the same covariates used in analyses of stimulant and sleep effects. The FC matrix for cetirizine is edge-for-edge correlated with that of stimulants at *r* = 0.10. Total *n* = 5,795, 291 taking cetirizine. For names and locations of networks see Supplemental Figure 1.

**Figure 8:**
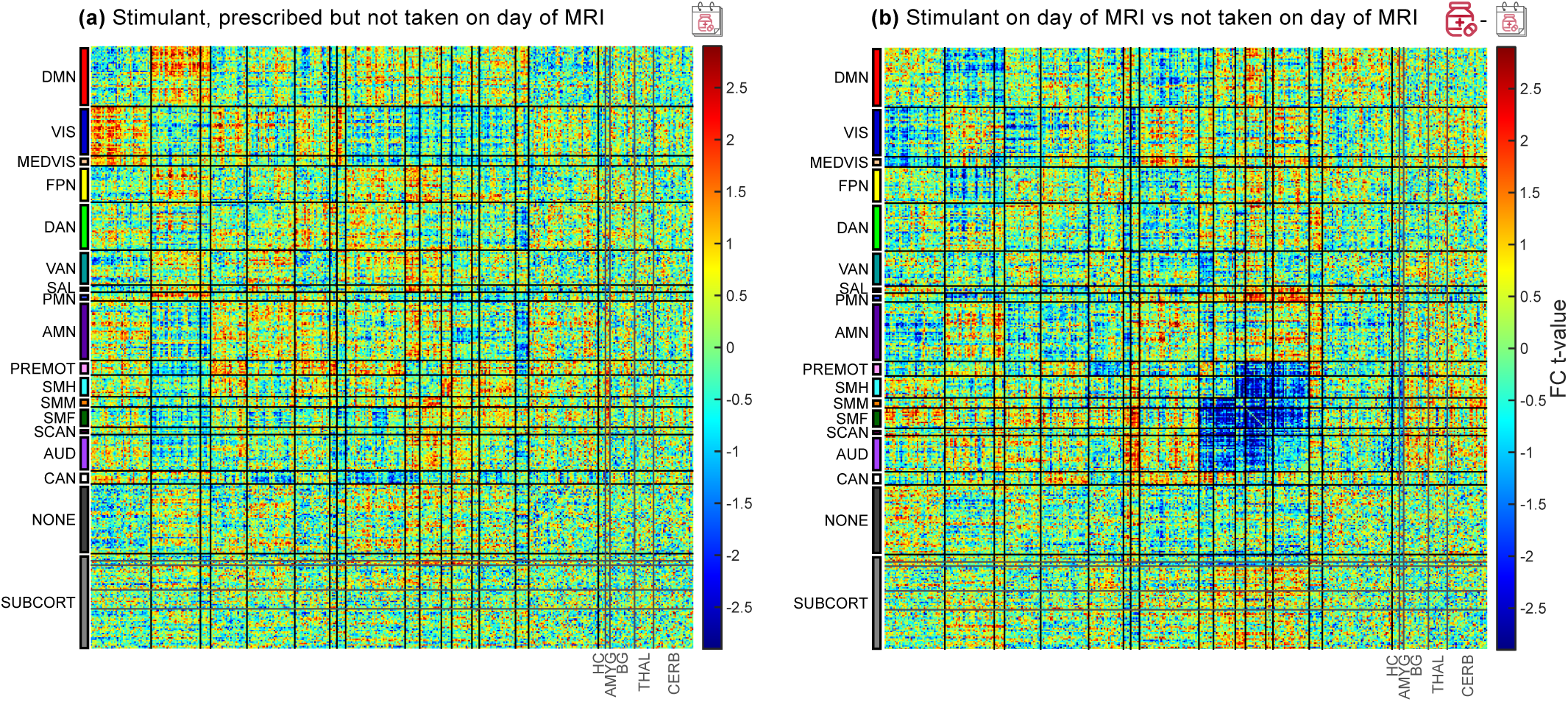
Differences in FC related to stimulants not taken on the day of scanning. **(a)** Difference between *n* = 76 children who were prescribed a stimulant but did not take it on the day of scanning and *n* = 5,382 children not prescribed a stimulant. The FC matrix was edge-for-edge correlated with that of stimulants taken on the day of scanning vs all other participants (Supplemental Figure 2) at *r* = 0.015. **(b)** Difference between *n* = 337 children who took a stimulant on the day of scanning and *n* = 76 children prescribed a stimulant who did not take it on the day of scanning. The FC matrix was edge-for-edge correlated with that of stimulants taken on the day of scanning vs all other participants at *r* = 0.55. For names and locations of networks see Supplemental Figure 1.

**Figure 9:**
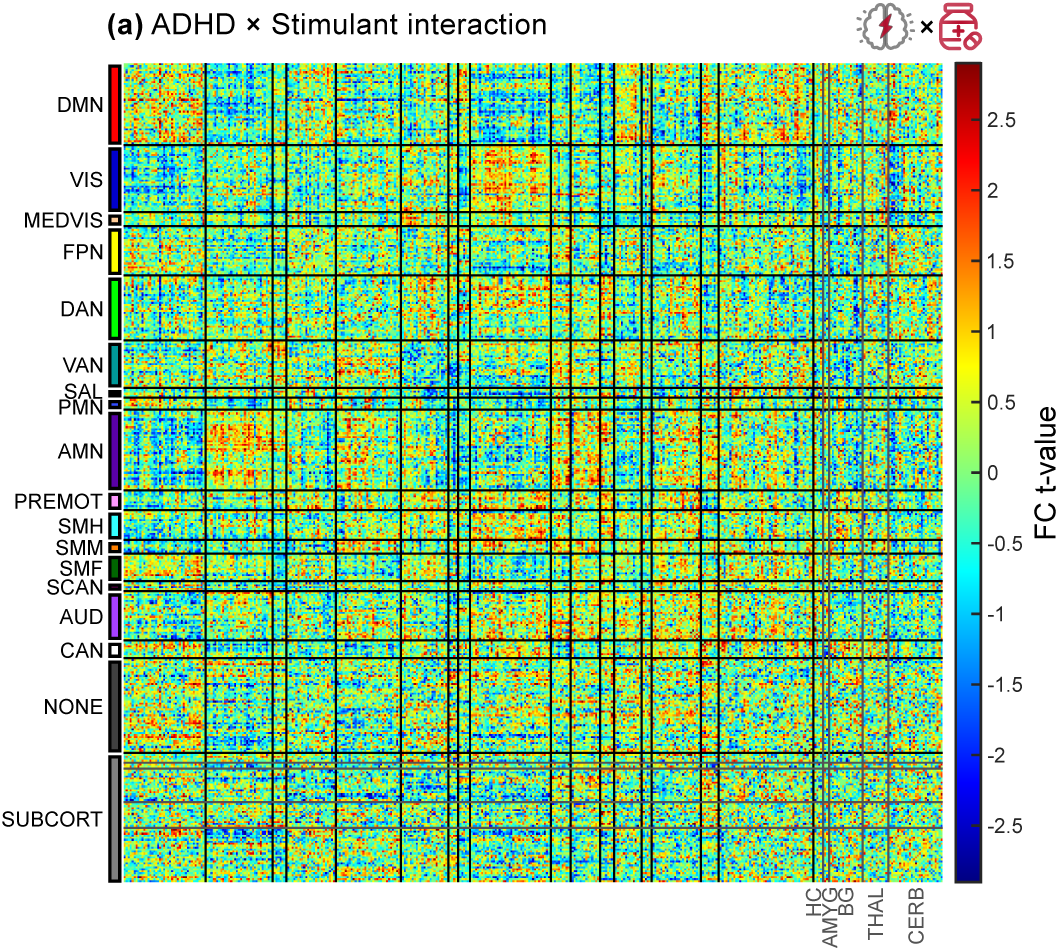
ADHD-specific difference in FC related to stimulants. Data are shown for an edgewise, linear model of stimulant × ADHD interaction with sex, ADHD, and stimulant as covariates. Total *n* = 5,795, 337 taking stimulants, 195 with ADHD, 67 with ADHD taking stimulants. For names and locations of networks see Supplemental Figure 1.

**Figure 10:**
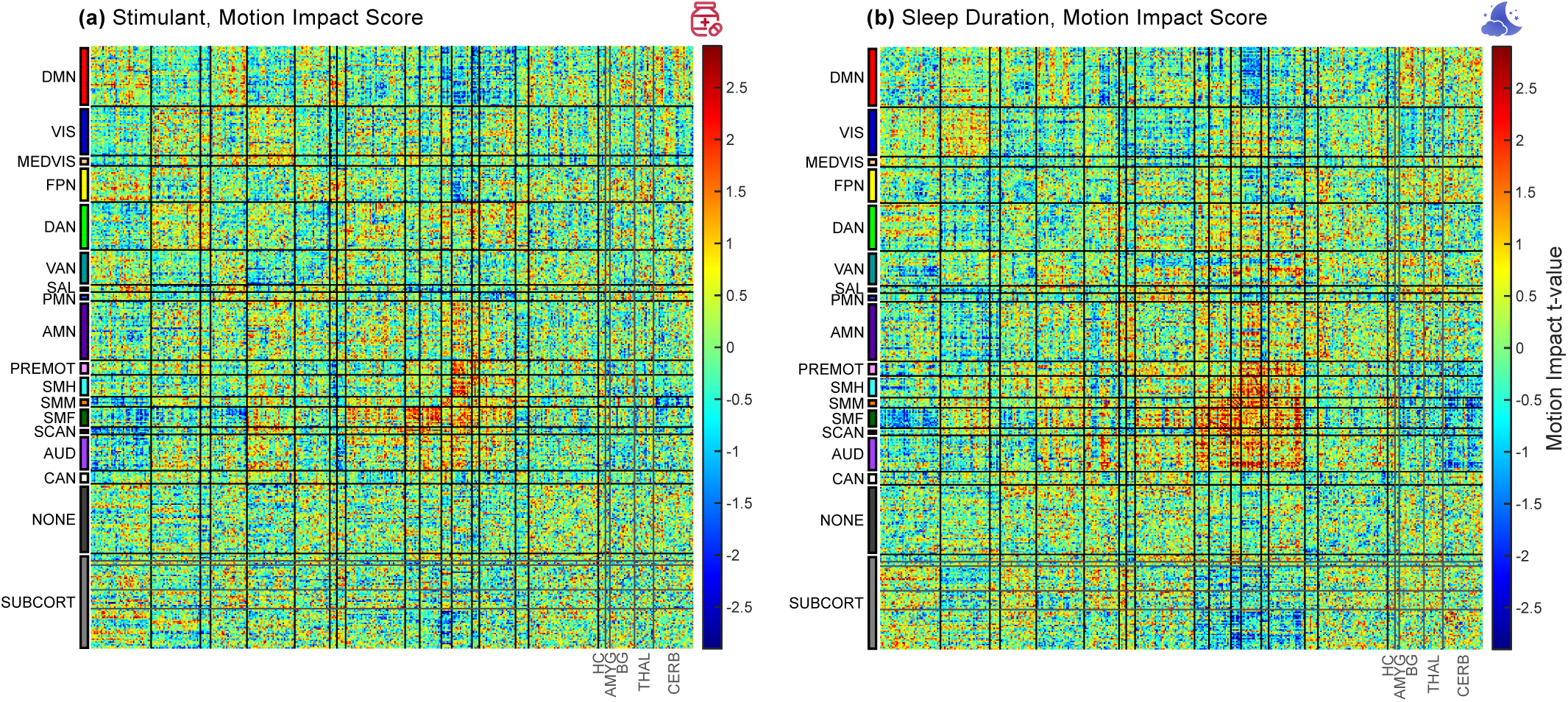
Motion Impact Assessment. Data were motion censored at FD < 0.2 mm. Motion impact scores reveal the effect of residual head motion artifact on stimulant- and sleep-related FC differences.^184^ Motion impact scores were anticorrelated with stimulant- and sleep-related FC differences, therefore the risk of motion-induced spurious findings is low. **(a)** Motion impact score for stimulants (*n* = 5,795, 337 taking stimulants). **(b)** Motion impact score for sleep duration (*n* = 5,795). See also Table 3. For names and locations of networks see Supplemental Figure 1.

**Figure 11:**
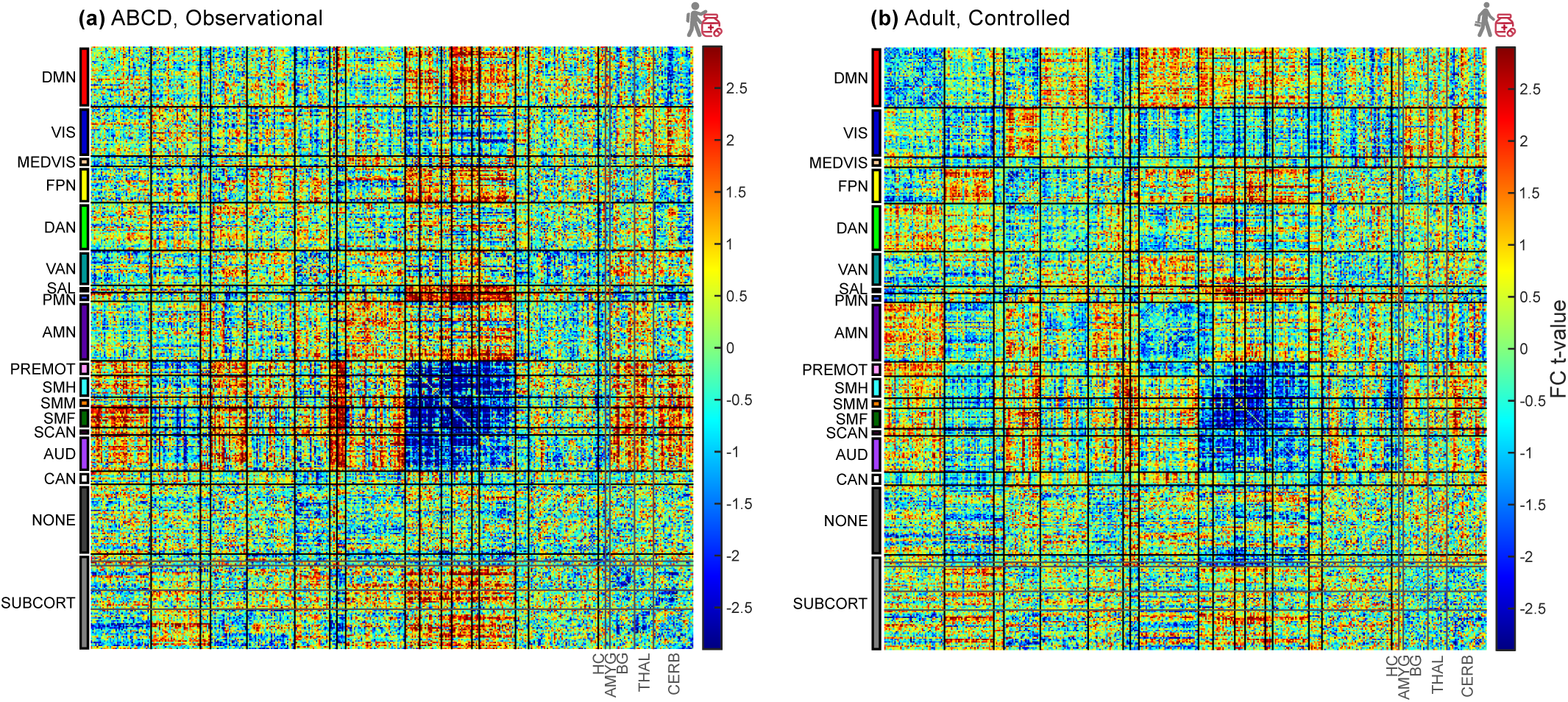
Comparison of stimulant-related FC differences across studies. **(a)** Children in the ABCD Study (*n* = 337 taking stimulant, *n* = 5,795 total). **(b)** Adults without ADHD in a controlled methylphenidate drug trial (*n* = 5).^123^ The FC matrices are edge-for-edge correlated at *r* = 0.18. For names and locations of networks see Supplemental Figure 1.

**Figure 12:**
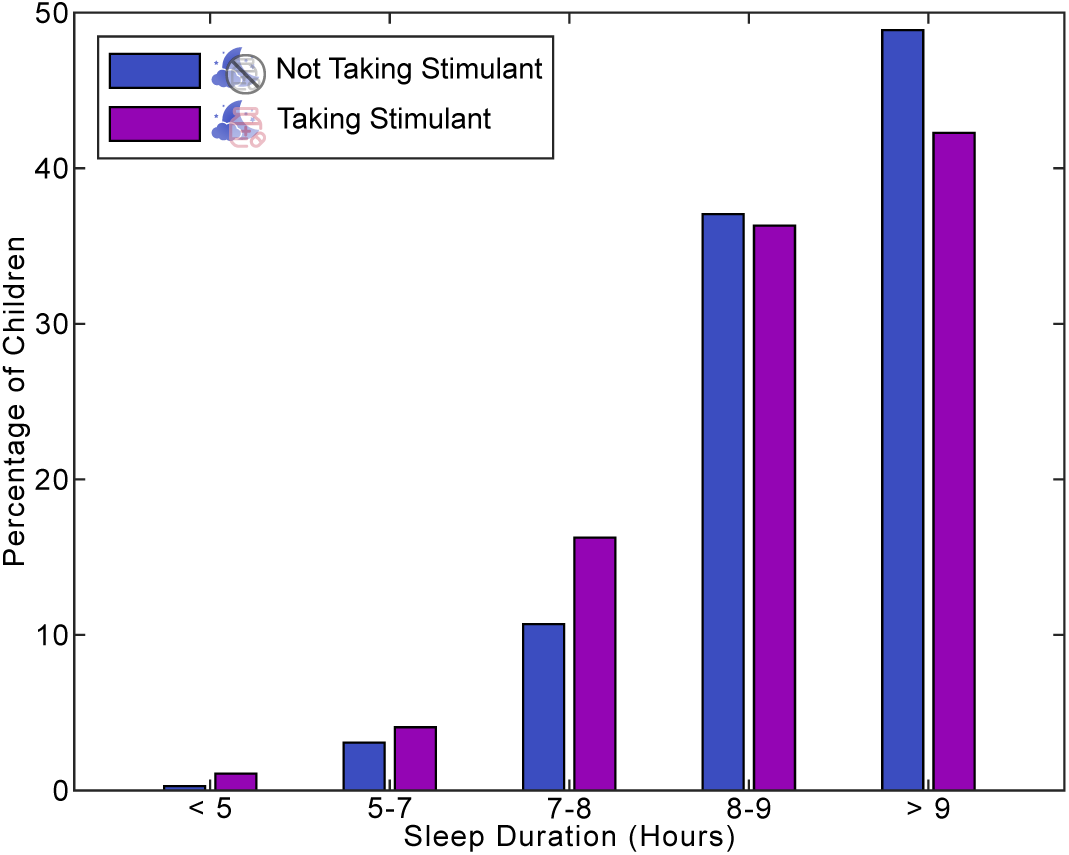
Sleep duration (in hours). Parent-reported average sleep duration is shown for children who did and did not take a stimulant on the day of scanning. For the purpose of reporting effect sizes we treated one ordinal unit of sleep duration as approximately equal to one hour (60 minutes) of sleep. Children who took a stimulant on the day of scanning got 10 fewer minutes of sleep per night (*n* = 5,795, 337 on stimulants, *P* = 1.0× 10^-4^). After controlling for ADHD (tier 4 criteria)^124^ the effect shrank to 7.3 minutes (*P* = 0.0068), and using less stringent criteria for ADHD the effect shrank to 1.7 fewer minutes of sleep per night (*P* = 0.54). ADHD diagnosis was associated with 20 fewer minutes of sleep per night (*n* = 175 with ADHD, *P* = 2.8× 10^-8^) by tier 4 criteria, or 15 minutes (*n* = 1142 with ADHD, *P* = 2.7×10^-19^) by less stringent criteria.

**Figure 13:**
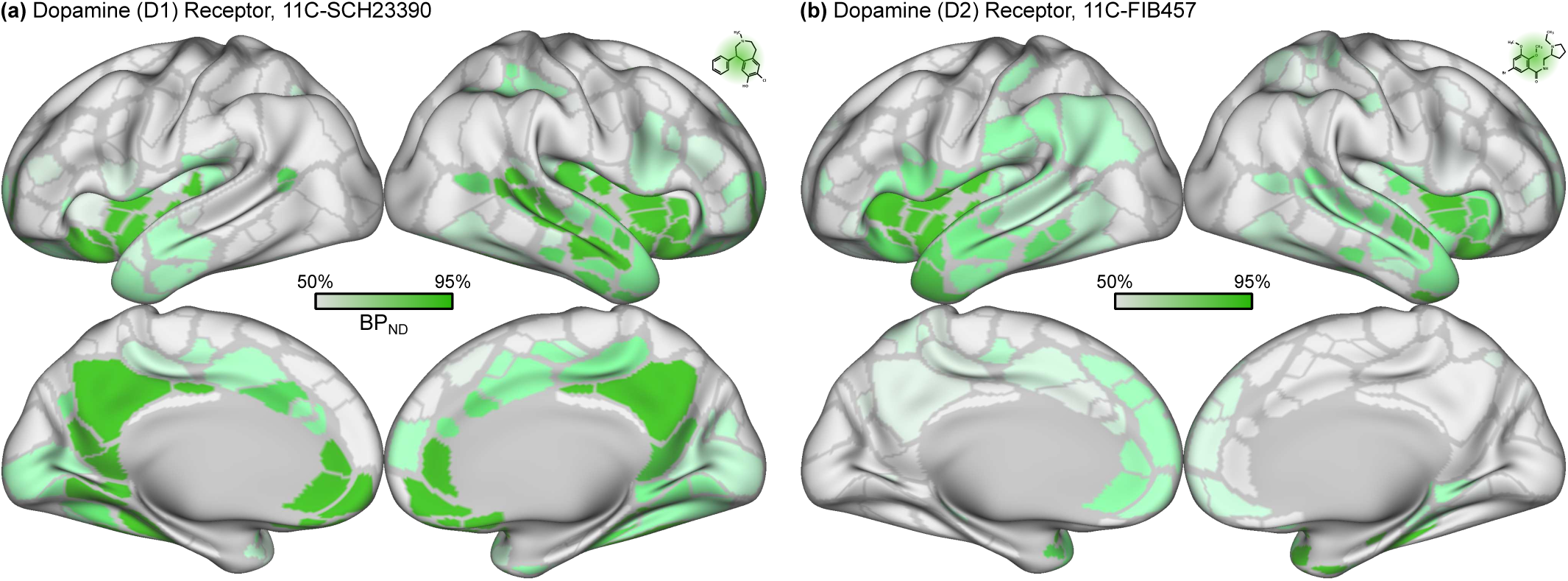
Dopamine receptor maps. Parcellated cortical receptor densities were obtained from positron emission tomography (PET) studies.^138^ D1 receptor maps were generated using the 11C-SCH23390 ligand (*n* = 13).^194^ D2 receptor maps were generated using the 11C-FLB457 ligand (*n* = 6).^195^ See also: Table 5.

**Figure 14:**
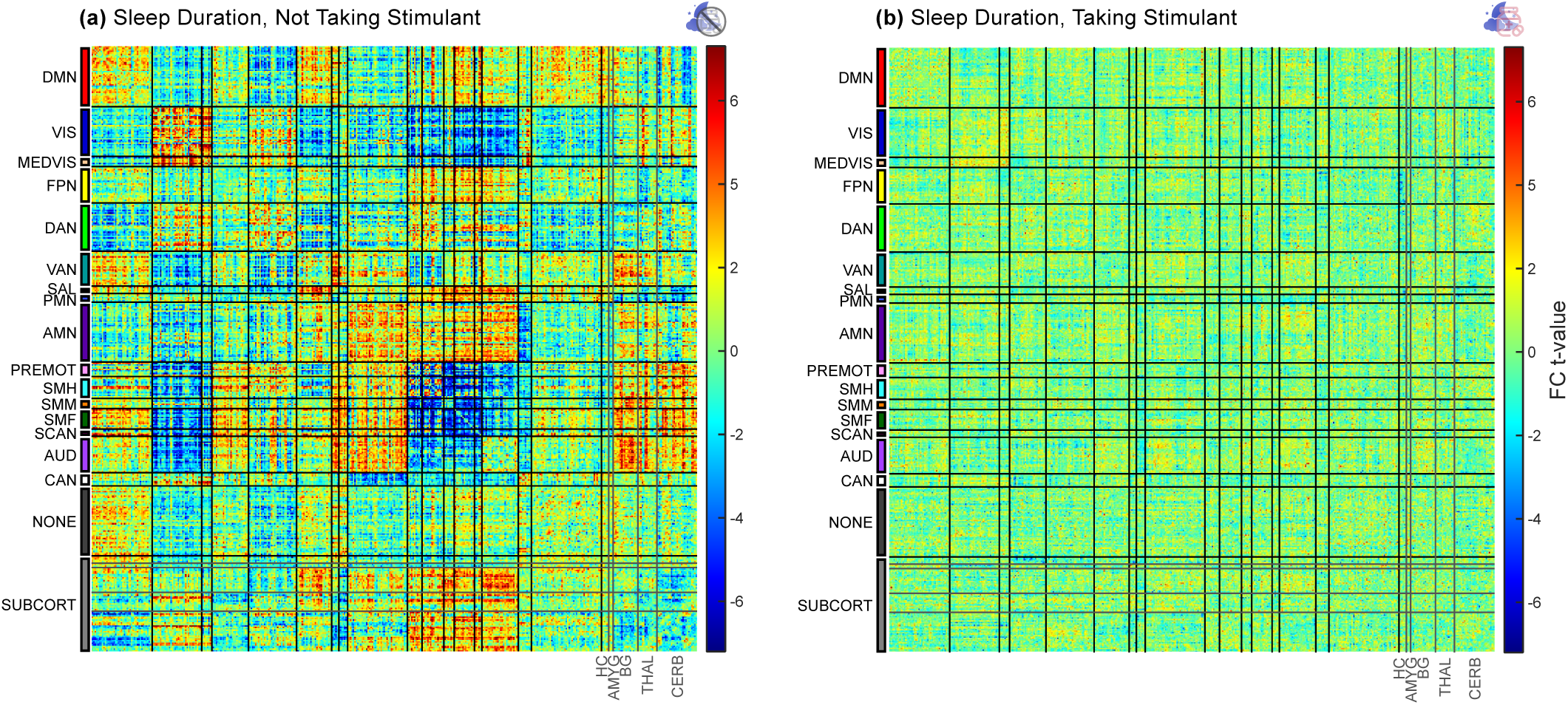
Differences in FC related to sleep in children on and off stimulants. **(a)** Children not taking a stimulant (*n* = 5,458). **(b)** Children taking a stimulant on the day of scanning (*n* = 337). For names and locations of networks see Supplemental Figure 1.

**Figure 15:**
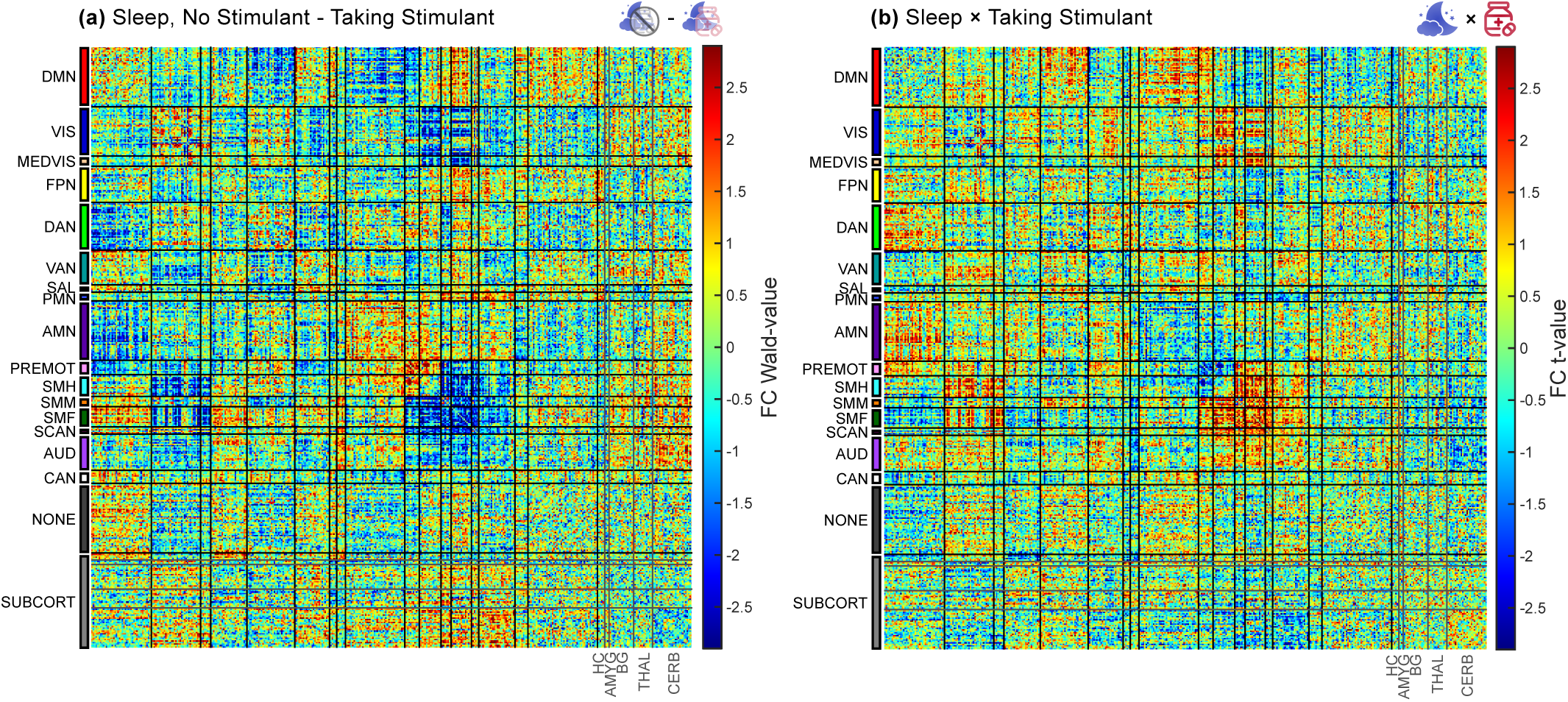
Comparison of differences in FC related to sleep in children on and off stimulants. **(a)** Wald test, sleep in children not taking stimulants (*n* = 5,458) minus sleep in children taking stimulants (*n* = 337). **(b)** Sleep × stimulant interaction (*n* = 5,795). For names and locations of networks see Supplemental Figure 1.

**Figure 16:**
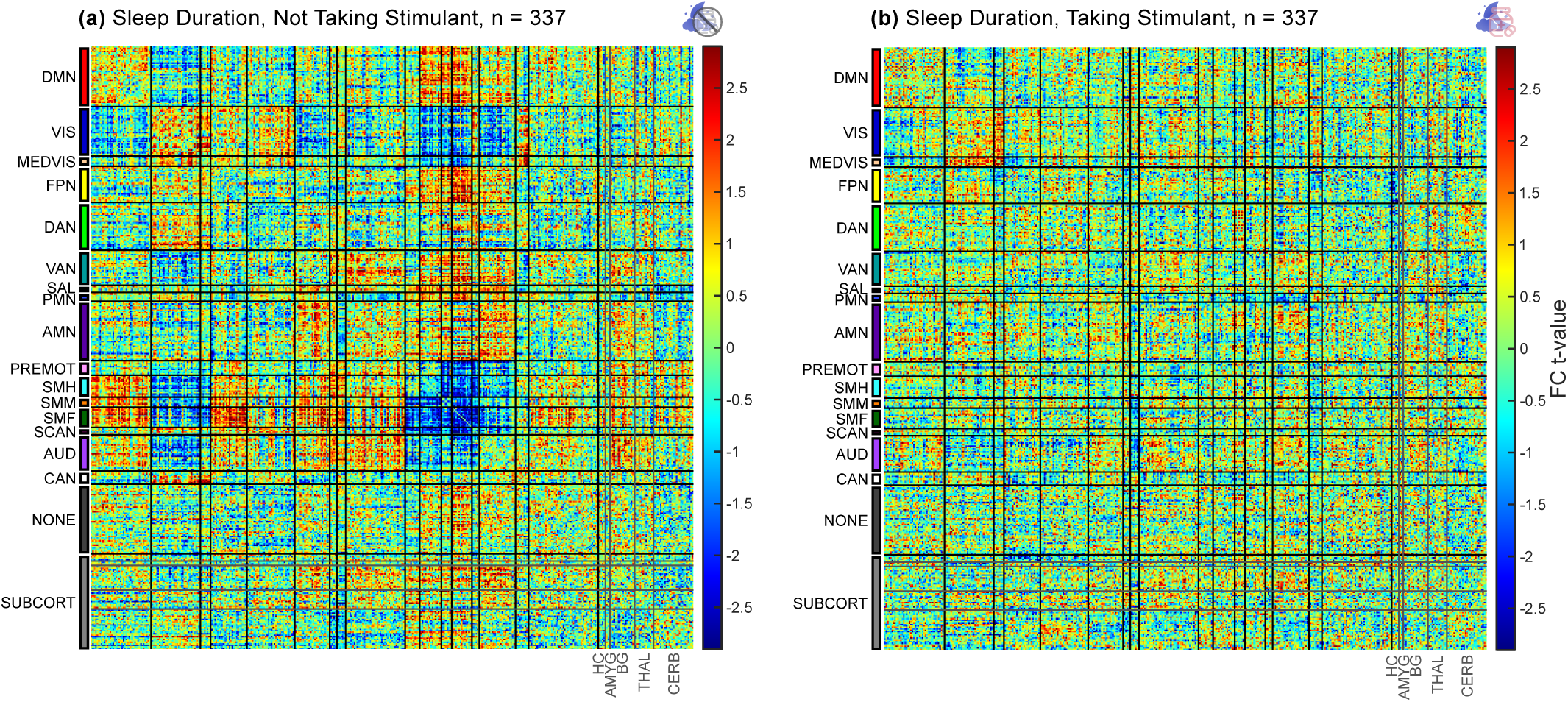
Differences in FC related to sleep matched for sample size. **(a)** Children not taking a stimulant (matched *n* = 337). **(b)** Children taking a stimulant (*n* = 337). For names and locations of networks see Supplemental Figure 1.

**Figure 17:**
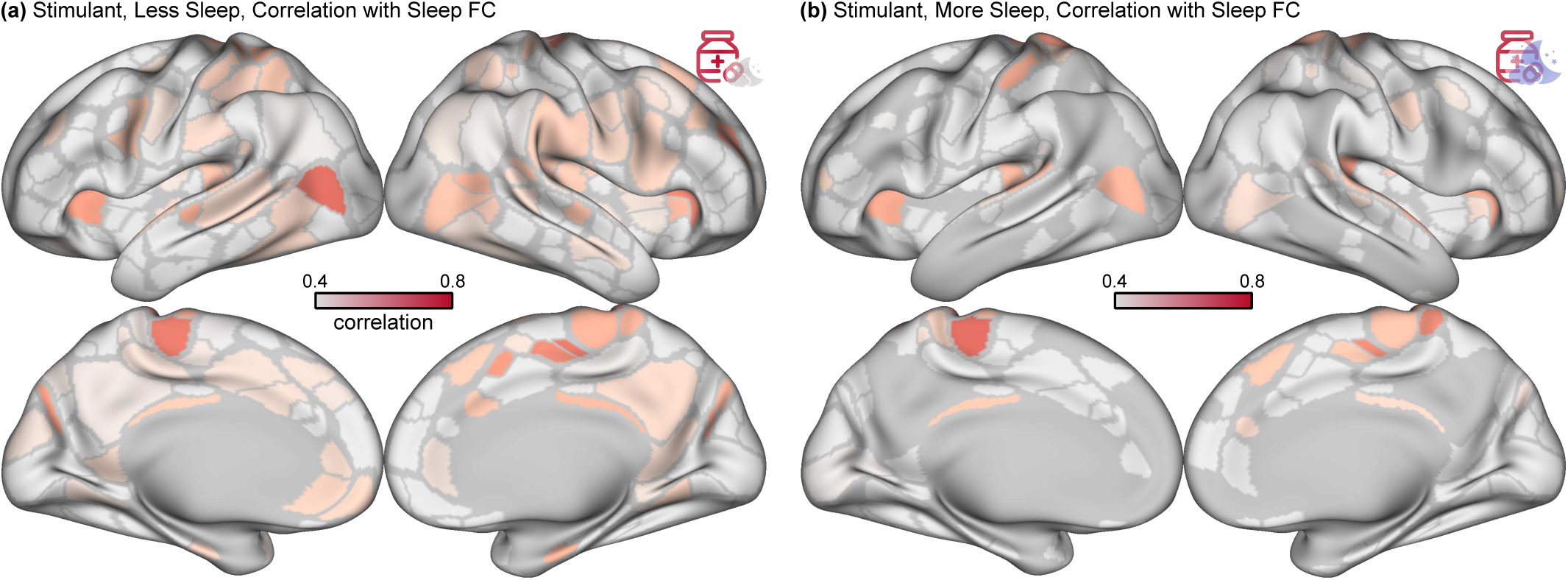
Parcelwise similarity of stimulant FC to sleep FC. FC differences related to stimulants were compared to FC differences related to sleep. The edgewise correlation between FC differences related to stimulants and sleep is shown for each cortical parcel. Negative values of correlation are shown in gray. **(a)** Children with less than 8 hours of sleep (*n* = 804, 68 takings stimulants). **(b)** Children with more than 8 hours of sleep (*n* = 2,883, 148 taking stimulants). The FC differences related to stimulants and sleep were more similar in children getting less sleep. For names and locations of networks see Supplemental Figure 1.

**Figure 18:**
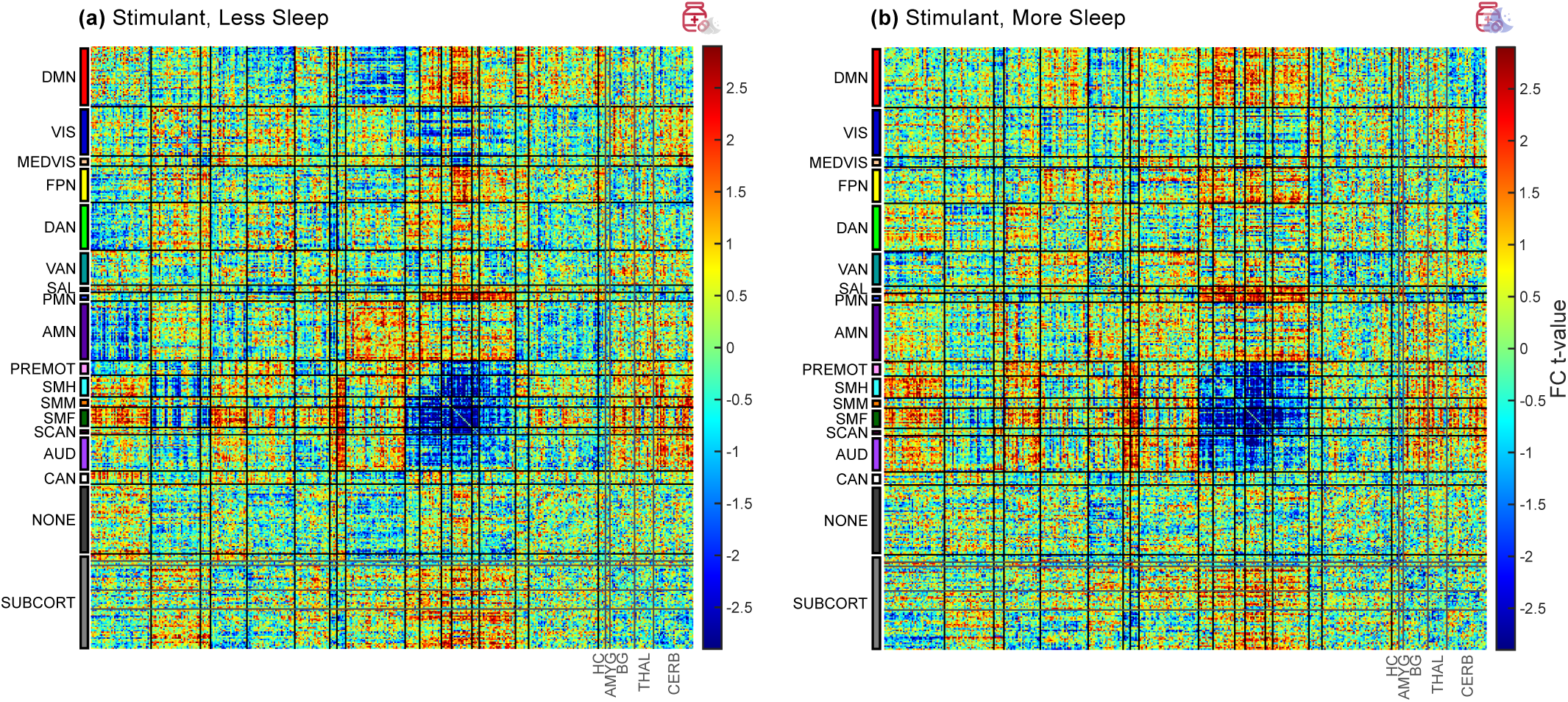
Differences in FC related to stimulants in children getting more or less sleep. **(a)** Children getting less than 8 hours of sleep (*n* = 804, 68 takings stimulants) and **(b)** children getting more than 8 hours of sleep (*n* = 2,883, 148 taking stimulants). For names and locations of networks see Supplemental Figure 1.

**Table 2:** Power analyses. Minimum detectable effect size (Cohen’s *d* ) at different power levels for stimulant-related differences in functional connectivity (FC) within attention and control networks. Effect sizes are for statistical inference with network level analysis (NLA). Previously reported effect sizes for stimulant-related FC differences in attention networks are approximately *d* = 0.89.^45^ NLA was at least 95% powered to detect FC differences of this size. DAN: dorsal attention network, VAN: ventral attention network, FPN: frontoparietal network.

|  | Effect Size |  |  |
| --- | --- | --- | --- |
| Power | DAN | VAN | FPN |
| 95% | 0.13 | 0.17 | 0.17 |
| 80% | 0.10 | 0.14 | 0.14 |

**Table 3:** Motion impact scores. An assessment of residual head motion artifact^184^ after motion censoring at FD < 0.2 mm was performed for stimulants and sleep duration. Significant motion overestimation scores indicate a risk of detecting spurious FC differences. Significant motion underestimation scores indicate a risk of failing to detect FC differences. Both stimulants and sleep had not-significant motion overestimation scores (*P* > 0.05, uncorrected), indicating an acceptably low risk of spurious findings due to head motion. Sample size *n* = 5,795. See also Supplemental Figure 10.

|  | Overestimation |  | Underestimation |  |
| --- | --- | --- | --- | --- |
| Variable | Z-score | $P$-value | Z-score | $P$-value |
| Stimulant | -4.97 | 0.890 | 10.5 | 0.085 |
| Sleep | -13.4 | 0.963 | 30.0 | 0.006 |

**Table 4:** Similarity of sleep and arousal brain maps. The ABCD Study included *n* = 5,795 children, 337 taking a stimulant. The norepinephrine transporter binding map was derived from positron emission tomography (PET) data using 11C-MRB (methylreboxetine) in *n* = 20 participants.^137,138^ The EEG alpha/delta power ratio (alpha slow wave index) map was derived from *n* = 10 participants.^117,118^ The respiratory variation map was generated using data from *n* = 190 participants from the Human Connectome Project.^113^ Correlations are reported for 333 cortical parcels.^133^ Significance testing was performed using spin tests.^129,130^ See also Figure 4.

|  | Stimulant |  | Sleep Duration |  |
| --- | --- | --- | --- | --- |
| | Correlation | $P$-value | Correlation | $P$-value |
| Norepinephrine | 0.36 | 0.0125 | 0.32 | 0.005 |
| EEG Alpha/Delta | 0.37 | 0.0085 | 0.49 | $< 0.0001$ |
| Respiratory Variation | 0.41 | 0.0225 | 0.51 | 0.0015 |

**Table 5:**
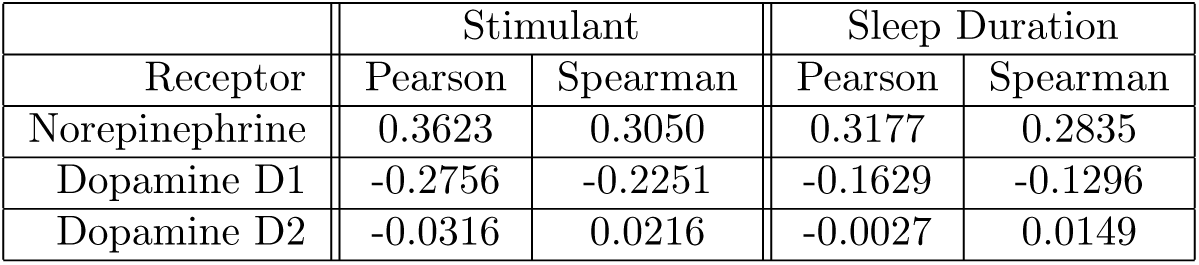
Correlation of FC differences with monoaminergic receptor densities. Parcellated cortical receptor densities were obtained from positron emission tomography (PET) studies,^138^ see Figure 13. Stimulant- and sleep-related differences in FC were strongly correlated with norepinephrine transporter density and weakly correlated with dopamine receptor density. Norepinephrine transporter (NET) receptors maps were generated using the 11C-MRB ligand (*n* = 20).^137^ D1 receptor maps were generated using the 11C-SCH23390 ligand (*n* = 13).^194^ D2 receptor maps were generated using the 11C-FLB457 ligand (*n* = 6).^195^ See also Figure 4 and Supplemental Figure 13.

|  | Stimulant |  | Sleep Duration |  |
| --- | --- | --- | --- | --- |
| Receptor | Pearson | Spearman | Pearson | Spearman |
| Norepinephrine | 0.3623 | 0.3050 | 0.3177 | 0.2835 |
| Dopamine D1 | -0.2756 | -0.2251 | -0.1629 | -0.1296 |
| Dopamine D2 | -0.0316 | 0.0216 | -0.0027 | 0.0149 |

## Supplemental Discussion

### Network level analysis facilitates understanding of FC

There has been much discussion in the literature about the statistical rigor of fMRI.^90^ FC matrices derived from the Gordon-Laumann-Seitzmann parcellation contain (394^2^-394)/2 = 77,421 distinct edges^133,180^ that make correction for multiple comparisons challenging. For example, Bonferroni correction for 77,421 comparisons would set a p-value threshold of 0.05/77,421 = 6.5× 10^-7^, greatly limiting statistical power.

Region of interest (ROI) analysis is one common strategy for reducing the number of statistical comparisons by using *a priori* knowledge to constrain which edges are investigated. Increased power to detect differences within the ROIs comes at the expense of decreased power to detect differences outside the ROIs. Among prior stimulant studies that selected ROIs in attention or cognitive control networks (DAN, VAN, or FPN), most^43,44,46–48,64^ found evidence supporting their *a priori* hypotheses of stimulant mechanism. The largest controlled (*n* = 99) ROI study found “surprisingly” no stimulant-related differences in attention networks,^60^ and the largest observational ROI study using ABCD data also did not detect significant results.^61^ None of these studies discussed stimulant-related differences in SM, SAL, or PMN, networks excluded by their ROIs. There are concerns that ROI analysis is susceptible to ascertainment bias within the ROIs (especially in small studies) and false negative results outside the ROIs.^96–98^

Network level analysis (NLA) reduces the number of statistical comparisons by imposing *a posteriori* constraints based on the network organization of the FC matrix without excluding any edges or regions.^121,122^ Our use of NLA allowed us to detect significant stimulant-related differences in action/motor networks and SAL/PMN even though we would not necessarily have predicted these changes based on a review of prior studies. Furthermore, visualizing the entire FC matrix allowed us to appreciate the relative magnitude of FC differences in SM, SAL, and PMN relative to attention networks (e.g. DAN). While ROI analysis remains a valid statistical approach, we anticipate that NLA and other data-driven whole-connectome approaches will enhance understanding of neurological phenomena involving multiple network interactions.

### Data collection in the ABCD Study

The ABCD Study was designed to prioritize breadth of information over detail.^119^ We were unable to model quantitative stimulant dose effects or learn the exact time during the day of scanning at which children took their stimulants. Many measures, including ADHD diagnosis, were derived primarily through parent report. As many as 79.3% of children prescribed a stimulant by parent report did not meet stringent criteria for ADHD, raising the question of why they were prescribed a stimulant. Without access to the prescribing physician’s medical record it is impossible to verify if reported prescriptions and ADHD criteria were accurate.

Data about sleep duration were also obtained by parent report. Even supposing these data were perfectly accurate, they only provide information about average sleep duration over many nights, and not sleep duration on the night before scanning specifically. Subsequent “waves” of the ABCD Study will collect accelerometry data (although only on a subset of children) that might provide a more accurate assessment of sleep duration on the night before scanning.

### Stimulant use and ADHD prevalence

Of the approximately 11,875 children recruited into the ABCD Study, several thousand children did not have usable imaging data or declined to completely answer questions about ADHD diagnosis, stimulant use, and other covariates leaving us with 5,795 children in this study. The exclusion of nearly half of prospective participants may have introduced sampling bias and inflated our estimates of ADHD and stimulant prevalence. Prior studies of the ABCD cohort have estimated the prevalence of ADHD to be anywhere from 3.5%^124^ to 10.3%^196^ depending on how the diagnosis was inferred from parent/child responses and how many children were included in the total count of participants. Prior estimates of stimulant use prevalence in children range from 5.3%^4^ to 6.1%^5^. A prior estimate of stimulant use from the ABCD data of 1.3% may have underestimated the true population prevalence by excluding children without a definite ADHD diagnosis.^196^

### Subcortical observations

Stimulants are widely believed to exert their dopaminergic effects through connections between the nucleus accumbens^30,78^ and cortical regions in the SAL.^53,68^ Evidence for this model in our data are equivocal. We observed large stimulant-related FC changes in SAL/PMN, but we we did not find significant differences in nucleus accumbens FC related to stimulants, see Supplemental Figure 3. Our ability to detect FC changes in basal ganglia and thalamus may have been affected by limitations of the NLA approach, which is biased to detect significant changes in physically larger brain regions,^121,122^ and high inter-individual variability in subcortical brain organization.^197^ Future fMRI studies using pulse-sequence protocols optimized for sensitivitiy to subcortical BOLD signal, highly-sampled individuals,^81,198^ and novel statistical methods are needed to elucidate the mechanisms of stimulant action in the subcortex.

